# mGluR6 coordinates cone terminal targeting and synaptic layer assembly during human retinal development

**DOI:** 10.64898/2026.04.23.720451

**Authors:** Leila Bahmani, Patricia Galvan, Hirad Hosseini, Jinlun Bai, Sumitha P. Bharathan, Michael Chen, Kayla Stepanian, Matthew E. Thornton, Brendan H. Grubbs, Aaron Nagiel

**Affiliations:** Vision Center, Department of Surgery, Children’s Hospital Los Angeles, Los Angeles, CA 90027, USA; Saban Research Institute, Children’s Hospital Los Angeles, Los Angeles, CA 90027, USA; Keck School of Medicine, University of Southern California, Los Angeles, CA 90033, USA; Maternal-Fetal Medicine Division, Department of Obstetrics and Gynecology, Keck School of Medicine, University of Southern California, Los Angeles, CA 90033, USA; Roski Eye Institute, Department of Ophthalmology, Keck School of Medicine, University of Southern California, Los Angeles, CA 90033, USA

**Keywords:** mGluR6, *GRM6*, inducible knockout, human retinal organoids, synaptic assembly, cone photoreceptor, synaptogenesis, outer plexiform layer, CRISPR/Cas9, Cre/loxP

## Abstract

The metabotropic glutamate receptor 6 (mGluR6), encoded by *GRM6*, is a core component of the ON-bipolar signaling cascade in the retina, but its role in human retinal development remains unclear. Here, we used temporally controlled CRISPR-based genetic ablation in human induced pluripotent stem cell–derived retinal organoids to define the developmental functions of mGluR6. Unexpectedly, we found that mGluR6 is expressed not only in depolarizing ON-bipolar cells but also transiently in cone photoreceptors during human retinal development, a pattern not observed in the mouse retina. Early loss of *GRM6* prior to synaptogenesis disrupted cone pedicle architecture, leading to mislocalization of synaptic proteins including Bassoon, ELFN2, and TRPM1, and ultimately resulting in widening or duplication of the outer plexiform layer (OPL). In contrast, deletion after synapse formation did not alter OPL synapses or morphology, revealing a temporally restricted requirement for mGluR6 during circuit assembly. These findings uncover a previously unrecognized role for mGluR6 in coordinating cone terminal targeting and synaptic layer assembly during human retinal development and highlight the power of temporally controlled genetic manipulation in organoid systems to reveal species-specific mechanisms of neural circuit formation.

## INTRODUCTION

Faithful visual perception depends on precisely organized synaptic transmission within the retina, where photoreceptors convert light into graded glutamate release that is interpreted by parallel ON and OFF pathways.^1,2^ At the first synapse of the visual system, ON-bipolar cells (ON-BCs) detect reductions in glutamate through the metabotropic glutamate receptor 6 (mGluR6), encoded by *GRM6*.^3^ In the mouse retina, mGluR6 is concentrated at ON-bipolar dendritic tips in the outer plexiform layer (OPL), positioned to couple photoreceptor output to compartmentalized postsynaptic signaling.^4^

In the dark, glutamate released from photoreceptors activates mGluR6 on ON-BCs, triggering a Gα_o_-mediated signaling cascade that closes the TRPM1 cation channel and hyperpolarizes the cell. Light-induced reduction in glutamate release relieves this inhibition, allowing TRPM1 channels to open and depolarize the ON-BCs.^5,6^ The fidelity of this pathway depends on the precise alignment of presynaptic photoreceptor ribbon release sites with postsynaptic receptor and signaling assemblies within the OPL.^7^ Disruption of mGluR6 signaling causes congenital stationary night blindness (CSNB), underscoring its essential role in ON-pathway transmission.^8,9^ Despite a detailed understanding of its role in mature synaptic transmission,^10–12^ whether mGluR6 contributes to earlier developmental processes in retinal circuit assembly remains poorly defined.

mGluR6 loss in mice abolishes ON-BC electrical responses (no b-wave) and mislocalizes key postsynaptic components, including TRPM1, GPR179, and RGS11,^11,13–15^ establishing mGluR6 as a central organizer of postsynaptic architecture. However, because these findings derive from constitutive knockout models, it remains unclear whether these deficits arise from impaired synapse assembly during development or from disassembly or faulty maintenance after synapses are formed. Notably, AAV-mediated *Grm6* gene augmentation in postnatal day 15 (P15) knockout mice only partially restored synaptic protein expression and failed to rescue ON-BC electrical responses,^16^ suggesting that mGluR6 may be required during the initial stages of synaptogenesis. Furthermore, existing models rely exclusively on the rod-dominant mouse retina, which lacks the specialized cone-rich architecture of human macula.^17,18^ Such species-specific differences highlight the importance of human-based systems to define how mGluR6 contributes to retinal synapse development and organization.

To overcome these limitations, we employed human retinal organoids (HROs), which recapitulate the cellular architecture, synaptic organization, and developmental trajectory of the mid-gestation human retina.^19^ Although HROs lack regional specializations such as macula, they display relatively high cone-to-rod ratios and therefore provide a valuable platform for studying human-specific aspects of retinal circuit formation.^20,21^ To achieve precise temporal control of *GRM6* disruption, we developed a tamoxifen-inducible knockout (iKO) model using CRISPR/Cas9 genome editing in combination with CreERT2/loxP technology.^22^ This system enabled targeted *GRM6* gene inactivation at specific stages of retinal development, allowing us to dissect whether mGluR6 is required for early synaptic layer assembly, trans-synaptic organization, or maintenance of mature retinal circuitry.

## RESULTS

### Human-specific cone expression of *GRM6* is recapitulated in retinal organoids

To define species-specific patterns of *GRM6* expression, we first analyzed publicly available single-cell RNA sequencing (scRNA-seq) datasets from mouse and human retina.^23,24^ In the mouse retina, *Grm6* transcripts were detected exclusively in bipolar cells (Figure 1A), consistent with prior work.^3,25,26^ In contrast, human retinal datasets revealed *GRM6* expression not only in bipolar cells but also in cone photoreceptors (Figure 1B). Notably, cone expression emerged as early as fetal week (FW) 14. While *GRM6* levels remained robust in bipolar cells throughout development, expression in cones progressively declined after FW20 and was undetectable in the adult human retina.

**Figure 1.**
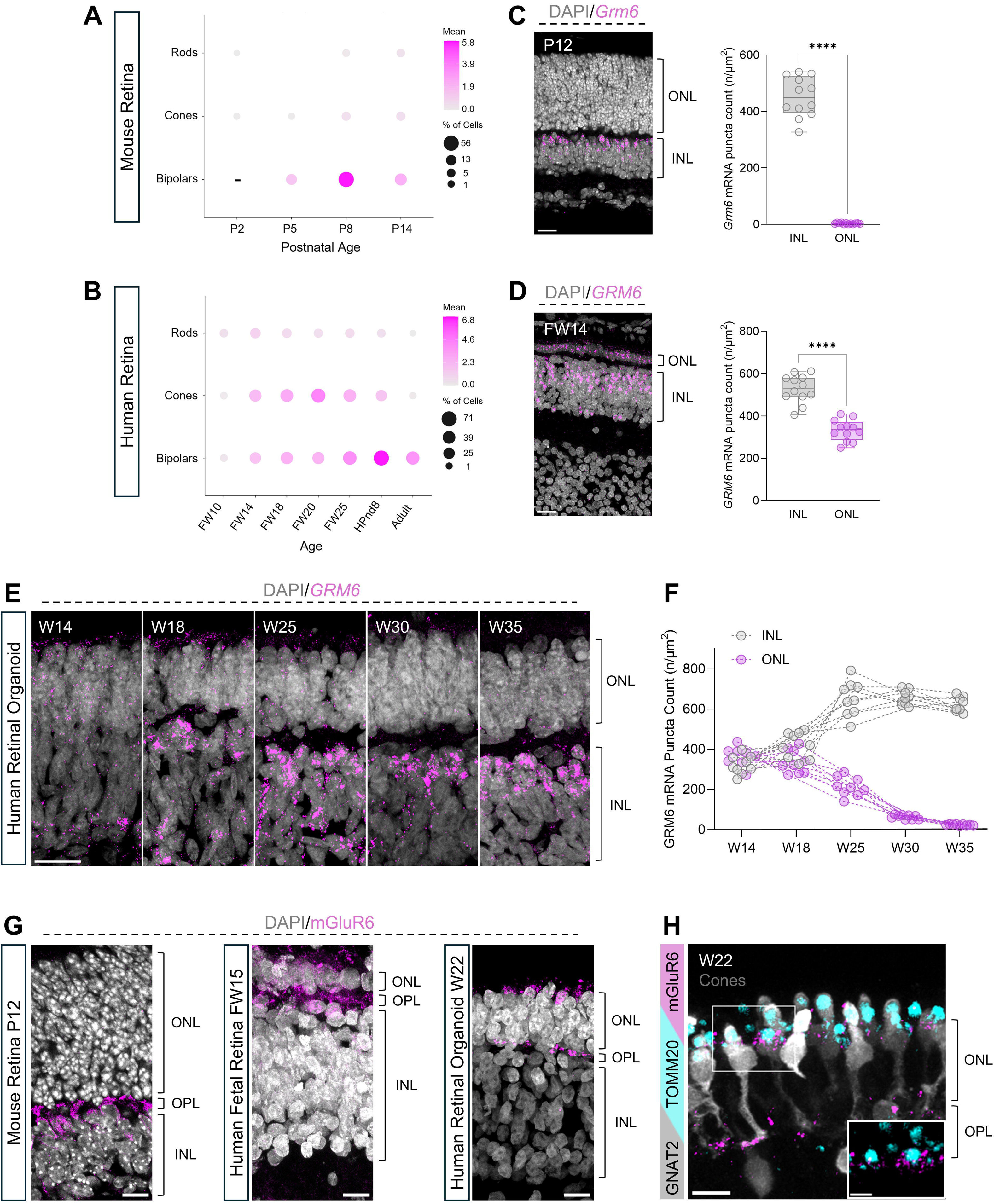
Human-specific cone expression of *GRM6* is recapitulated in retinal organoids. (A–B) Bubble plots of scRNA-seq datasets from mouse (A) and human (B) retina showing *GRM6* expression across developmental timepoints. Color intensity (gray to magenta) indicates the mean expression level, and bubble size represents the percentage of cells expressing *GRM6*. Absence of bipolar cells at early stages is indicated by “–”. (C–D) RNA-FISH detection of *GRM6* transcripts (magenta) in postnatal day 12 (P12) mouse retina (C) and fetal week 14 (FW14) human retina (D). Nuclei are counterstained with DAPI (gray). Scale bars, 20 μm. Right panels show quantification of diffraction-limited *GRM6* RNA puncta within the inner nuclear layer (INL) and outer nuclear layer (ONL). (E) Representative RNA-FISH images of human retinal organoids (HROs) showing *GRM6* transcript (magenta) distribution across developmental stages (W14-W35; W, weeks in culture). Nuclei are counterstained with DAPI (gray). Scale bar, 20 μm. (F) Quantification of diffraction-limited *GRM6* RNA puncta within INL and ONL of HRO sections across developmental stages. (G) Representative immunofluorescence images detecting mGluR6 protein (magenta) in mouse retina, human fetal retina, and HROs. OPL, outer plexiform layer. Nuclei are counterstained with DAPI (gray). Scale bar, 10 μm. (H) Representative immunofluorescent image of HROs co-labeled for mGluR6 (magenta), mitochondrial marker TOMM20 (cyan), and cone marker GNAT2 (gray). Scale bar, 10 μm; Inset 5 μm. Quantification was performed on 12 mouse retinal sections (P10–P12) and 12 human fetal retinal sections (FW14–FW16). For HROs, 9 organoids per stage from 3 independent differentiation batches were analyzed. Four regions of interest (ROIs) per section or organoid were quantified, and values represent the mean ± SD per sample. Comparisons between INL and ONL *GRM6* expression were performed using unpaired two-tailed t tests, ****P < 0.0001.

To validate these transcriptomic findings *in situ*, we performed RNA fluorescence *in situ* hybridization (RNA-FISH) on mouse and human retinal sections. In the mouse retina, *Grm6* RNA puncta were confined to the inner nuclear layer (INL), where bipolar cell somata reside (Figure 1C). Quantification of *Grm6* RNA puncta revealed abundant INL expression (451.3 ± 70.9 puncta per 250 μm²) with negligible signal in the outer nuclear layer (ONL; 2.8 ± 2.1 puncta), the layer containing rod and cone photoreceptor nuclei. In contrast, human fetal retina exhibited robust *GRM6* expression in both INL (528.2 ± 65.4 puncta) and ONL (332.4 ± 49.4 puncta) (Figure 1D), confirming cone-associated expression in mid-gestation human retina.

We next examined whether human retinal organoids recapitulate this developmental pattern. RNA-FISH across organoid maturation (W14–W35; W, weeks in culture) demonstrated *GRM6* puncta in both INL and ONL compartments (Figure 1E). Quantitative analysis revealed a progressive decline in ONL expression after W18 (Figure 1F), closely paralleling the temporal dynamics observed in human fetal retina.

To determine whether these transcriptional patterns were reflected at the protein level, we performed immunofluorescence staining for mGluR6 in mouse retina (P12), human fetal retina (FW15), and HROs (W22). In the mouse retina, mGluR6 protein is localized exclusively to the OPL, corresponding to the dendritic processes of ON-BCs (Figure 1G). In contrast, human fetal retina and HROs exhibited mGluR6 signal not only in the OPL but also within the ONL (Figure 1G). Notably, the ONL-associated mGluR6 signal appeared enriched toward the apical region of the layer. Using age-matched GNAT2 reporter HROs, in which cone photoreceptors express GFP,^27^ we confirmed that mGluR6 immunoreactivity is present within cone photoreceptors in addition to its canonical localization in the OPL (Figure 1H). Co-staining with the mitochondrial marker TOMM20, which labels photoreceptor inner segments, revealed that the mGluR6 signal localizes predominantly basal to the TOMM20-positive mitochondrial layer and apical to cone nuclei within the ONL. This subcellular distribution corresponds to the inner segment biosynthetic compartment, which is enriched in endoplasmic reticulum and Golgi apparatus,^28^ consistent with intracellular synthesis and processing of mGluR6 within cone photoreceptors (Figure 1H).

Together, these data identify a species-specific difference in *GRM6* expression, revealing transient cone photoreceptor expression in the developing human retina that is absent in mice and faithfully recapitulated in human retinal organoids. Building on this foundation, we next generated a tamoxifen-inducible *GRM6* knockout system in HROs to dissect the temporal roles of mGluR6 in human retinal development.

### Temporal control of *GRM6* knockout in human retinal organoids

To enable precise temporal control of *GRM6* ablation, we engineered a tamoxifen-inducible *GRM6* knockout (iKO) system in the human WTC-11 iPSC line. Two guide RNAs (gRNAs) were designed using the CRISPOR web tool^29^ to target sequences flanking exons 1 and 2 of *GRM6* (Figure 2A). These guides enabled homology-directed insertion of loxP sites through co-delivery of 134-bp single-stranded oligodeoxynucleotide (ssODN) donor templates. Following single-cell cloning, biallelic floxing of exons 1 and 2 was confirmed by PCR genotyping and Sanger sequencing (Figure S1A; Table S1).

**Figure 2.**
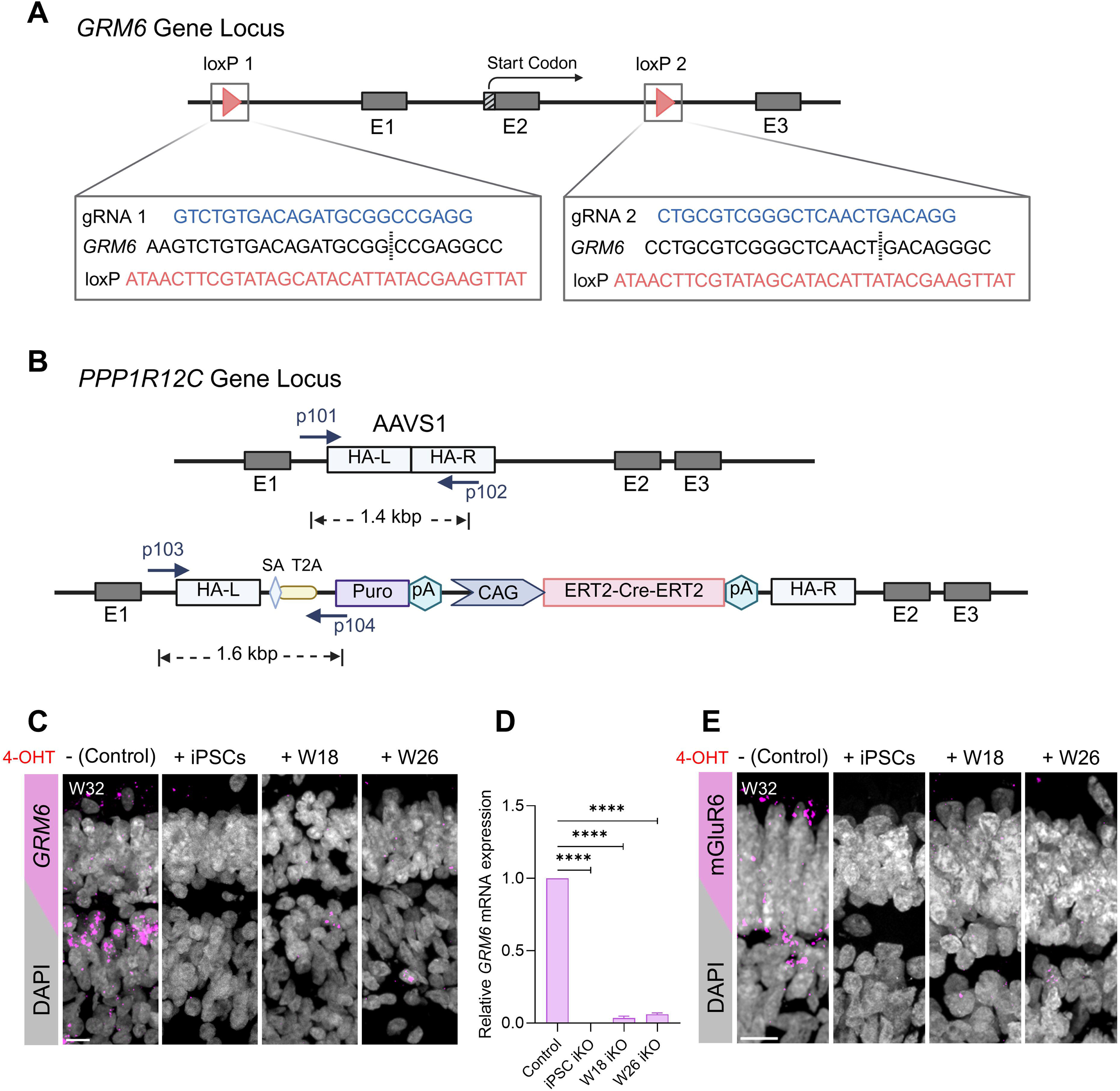
Temporal control of *GRM6* knockout in human retinal organoids. (A) Schematic of the *GRM6* gene locus showing CRISPR/Cas9-mediated insertion of loxP sites flanking exons 1 and 2. Vertical dashed lines indicate the Cas9 cleavage and *loxP* insertion sites. E, exon; gRNA, guide RNA. (B) Schematic of targeted integration of the ERT2–Cre–ERT2 cassette into the AAVS1 locus within the *PPP1R12C* gene. The donor construct includes homology arms (HA-L and HA-R), a CAG promoter–driven ERT2–Cre–ERT2 cassette, and a puromycin resistance gene (Puro). PCR genotyping strategy and expected amplicon sizes for edited and unedited alleles are indicated. (C) Representative RNA-FISH images showing *GRM6* transcripts (magenta) in week 32 (W32) vehicle-treated control and iKO HROs following 4-hydroxytamoxifen (4-OHT) induction at the indicated developmental stages. Scale bar, 10 μm. (D) Quantification of relative *GRM6* mRNA levels by quantitative PCR (****P < 0.0001, one-way ANOVA followed by Dunnett’s multiple-comparisons test). Data are presented as mean ± SD. Total RNA was extracted from 9 W32 HROs derived from 3 independent differentiation batches (n = 3 biological replicates per condition; 3 organoids per batch). (E) Representative immunofluorescence images indicating mGluR6 protein (magenta) in W32 vehicle-treated control and iKO HROs following 4-OHT induction at the indicated developmental stages. Nuclei are counterstained with DAPI (gray). Scale bar, 10 μm. See also Figures S1-S3 and Tables S1 and S2.

To confer inducible recombination, an ERT2-Cre-ERT2 cassette was inserted into the safe-harbor AAVS1 locus of the floxed iPSCs (Figure 2B). The dual-ERT2 configuration was selected to minimize leaky recombination and ensure stringent tamoxifen-dependent Cre activation.^30^ Correct monoallelic ERT2-Cre-ERT2 integration was verified by PCR (Figure S1B; Table S1). Quality control analyses confirmed that the resulting *GRM6*-iKO iPSC clones retained normal karyotype, genomic stability, and pluripotency marker expression, and exhibited no detectable off-target editing (Figures S1C–F; Tables S1 and S2), establishing their suitability for downstream differentiation.

We next assessed whether the edited iPSCs maintained retinal differentiation competence. Using a stepwise differentiation protocol,^19^ *GRM6*-iKO iPSCs formed embryoid bodies (day 1), neuroepithelial vesicles (day 15), and laminated HROs by day 45, with morphology and developmental timing indistinguishable from wild-type HROs (Figure S2A). In the absence of tamoxifen induction, RNA-FISH and immunofluorescence analyses at W25 confirmed robust *GRM6* transcript and mGluR6 protein expression, respectively, in both bipolar cells and photoreceptors of iKO organoids, comparable to wild-type HROs (Figures S2B and S2C). These findings indicate that retinal lineage specification and baseline *GRM6* expression remain intact prior to Cre activation.

To validate the functionality and temporal control of the inducible knockout system, either iKO iPSCs or HROs were treated with 4-hydroxytamoxifen (4-OHT) at defined developmental stages. For early ablation, iKO iPSCs were exposed to 2 μM 4-OHT for 48 hours, two days before retinal differentiation (iPSC iKO group). To target the onset of OPL synaptogenesis, which begins around W21, iKO HROs were treated with 4 μM 4-OHT for 7 days at W18 (W18 iKO group). To assess the role of mGluR6 in synapse maintenance, which reaches a plateau by W26, a third cohort received 4 μM 4-OHT at W26 (W26 iKO group). Vehicle-treated HROs from the same differentiation batch served as controls, and all groups were collected at W32 for subsequent analyses. Importantly, the W32 collection timepoint provides sufficient time to observe the consequences of mGluR6 depletion following W26 knockout. Although the half-life of mGluR6 has not been directly measured in the human retina, computational analyses using ProtParam^31^ and PLTNUM^32^ approaches indicate that mGluR6 is likely a relatively stable protein with a predicted half-life in the multi-day range (∼24–96 hours) in both human and mouse (Figure S3A). These estimates are consistent with experimentally measured half-lives of related Group III mGluRs, including mGluR4^33^ and mGluR7^34^, which also exhibit multi-day stability in neuronal tissues. Furthermore, mGluR6 is highly conserved between human and mouse (93.2% sequence identity)^35^ (Figure S3B), supporting similar turnover kinetics across species. Based on these estimates, the six-week interval between W26 knockout induction and W32 collection corresponds to approximately 10–42 protein half-lives, supporting effective depletion of existing protein and allowing sufficient time for downstream phenotypic effects to emerge.

RNA-FISH analysis revealed robust *GRM6* expression in vehicle-treated control HROs, whereas *GRM6* puncta were undetectable in iPSC iKO HROs and markedly reduced in both W18 and W26 iKO HROs (Figure 2C). Consistently, quantitative PCR analysis demonstrated complete loss of *GRM6* transcripts in iPSC iKO HROs and less than 7% residual expression in the W18 and W26 iKO groups compared to controls (Figure 2D). Immunofluorescence staining further confirmed efficient ablation at the protein level: mGluR6 immunoreactivity was completely absent in iPSC iKO HROs and reduced to negligible levels in W18 and W26 iKO HROs relative to controls (Figure 2E).

Together, these results establish a robust, tightly regulated inducible knockout platform that enables stage-specific ablation of mGluR6 during human retinal organoid development. This system provides the foundation to dissect the temporal roles of mGluR6 in retinal differentiation, synapse assembly, and circuit maintenance.

### Early *GRM6* deletion disrupts OPL laminar organization

To determine how temporally controlled *GRM6* deletion affects photoreceptor–bipolar synapse organization, we examined key components of the presynaptic ribbon, trans-synaptic complex, and postsynaptic signaling machinery (Figure 3A). The schematic in Figure 3A outlines the molecular architecture of the photoreceptor ribbon synapse, including presynaptic proteins (Bassoon, CtBP2/RIBEYE, PSD-95), trans-synaptic organizers (LRIT3, ELFN2), and postsynaptic ON-bipolar components (TRPM1, GPR179, RGS11), which guided our stage-specific structural analysis following inducible *GRM6* ablation.

**Figure 3.**
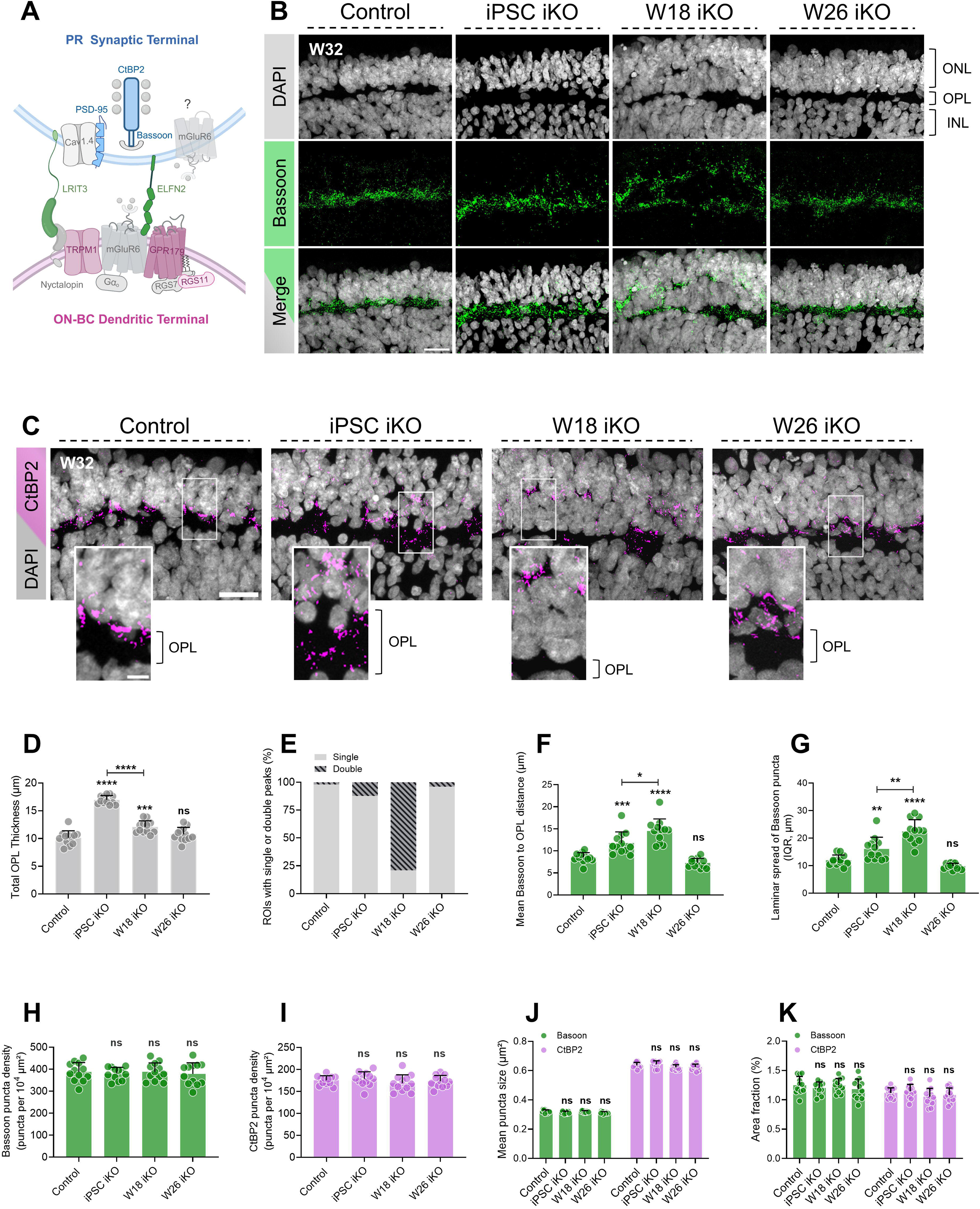
Early *GRM6* deletion disrupts OPL laminar organization. (A) Schematic representation of the photoreceptor (PR)–ON bipolar cell (ON-BC) synapse within the outer plexiform layer (OPL). (B-C) Representative immunofluorescence images of Bassoon (green; B) and CtBP2 (magenta; C) in week 32 (W32) inducible knockout (iKO) human retinal organoids (HROs) following stage-specific *GRM6* deletion. ONL, outer nuclear layer; INL, inner nuclear layer. Nuclei are counterstained with DAPI (gray). Scale bars, 20 µm (B) and 10 µm (C). (D) Quantification of total OPL thickness using vertical line profiling along the ONL–OPL–INL axis based on DAPI signal. (E) Percentage of regions of interest (ROIs) exhibiting single versus double Bassoon intensity peaks based on line profile analysis. (F–G) Quantification of Bassoon puncta positional distribution relative to the ONL-OPL boundary: (F) mean puncta distance and (G) interquartile range (IQR) of puncta distances. (H–K) Quantification of presynaptic protein parameters within standardized ROIs spanning ONL, OPL, and adjacent INL: (H) Bassoon puncta density, (I) CtBP2 puncta density, (J) mean puncta size, and (K) area fraction. Quantifications were performed on 12 organoids per group from 3 independent differentiation batches (n = organoids). For panels D and E, five measurements per ROI were obtained; for panels D–K, four ROIs per organoid were analyzed, and values represent the mean ± SD per organoid. Statistical analyses were performed using one-way ANOVA with Dunnett’s multiple-comparisons test relative to the control. Direct comparisons between iPSC and W18 groups were performed using unpaired two-tailed t tests. Exact P values are as follows: (D) ***P = 0.0003; (F) ***P = 0.0007, *P = 0.0176; and (G) **P = 0.0075, **P = 0.0013 (iPSC versus W18). ****P < 0.0001. See also Figures S4 and S5.

At W32, control organoids exhibited a single, sharply defined synaptic band in which Bassoon and CtBP2 puncta were confined to a compact OPL (Figures 3B and 3C). *GRM6* deletion at the iPSC stage resulted in a broadened OPL, with ribbon puncta dispersed within. In contrast, deletion at W18 led to the formation of discontinuous strips of pseudo-duplicated OPL. Finally, deletion at W26 showed no differences compared to the controls.

To quantify these structural changes, we performed standardized vertical line profiling across the ONL–OPL–INL axis (Figures S4). DAPI intensity profiles defined nuclear boundaries and total OPL thickness, measured as the distance between ONL and INL nuclear peaks (Figure S4A). In W18 iKO organoids, the combined thickness of both primary and pseudo-OPL bands was quantified as total OPL thickness. OPL thickness was significantly increased in iPSC iKO (16.9 ± 0.7 µm) and W18 iKO organoids (12.0 ± 1.1 µm) relative to controls (10.1 ± 1.2 µm), whereas W26 iKO organoids (10.7 ± 1.2 µm) were comparable to controls (Figure 3D).

Bassoon intensity profiles further resolved synaptic stratification (Figure S4B). Nearly all control and W26 iKO organoids exhibited a single dominant intensity peak (97.9% and 95.8%, respectively). In contrast, 79.1% of W18 iKO organoids displayed two spatially separated peaks, consistent with pseudo-duplication of the synaptic layer (Figure 3E). iPSC iKO organoids primarily exhibited broadened single peaks (87.5% single versus 12.5% double), consistent with diffuse expansion rather than discrete stratification (Figures 3E and S4B).

We next quantified ribbon positioning by calculating the mean distance and interquartile range (IQR) of Bassoon puncta relative to the ONL–OPL boundary (Figures 3F and 3G). Both metrics were significantly increased in iPSC and W18 iKO organoids compared to controls (mean distance: 11.7 ± 2.5 µm and 14.5 ± 2.7 µm versus 8.4 ± 1.1 µm; IQR: 16.1 ± 4.2 µm and 22.4 ± 4.2 µm versus 11.9 ± 1.9 µm). Importantly, these alterations were significantly greater in W18 iKO organoids than in iPSC iKO samples, supporting the emergence of two partially segregated ribbon strata rather than simple OPL broadening. W26 iKO organoids remained indistinguishable from controls (mean distance: 7.2 ± 1.0 µm; IQR: 9.8 ± 1.1 µm), indicating preserved laminar organization when *GRM6* was deleted after synaptic maturation.

Despite these pronounced spatial defects, presynaptic protein abundance was unchanged. Quantification within standardized regions of interest (ROIs) spanning the ONL, OPL, and adjacent INL revealed no significant differences in Bassoon or CtBP2 puncta density, mean puncta size, or area fraction, which reflects the percentage of ROI area occupied by signal (Figures 3H–K). Consistent with these findings, PSD-95, a presynaptic scaffold protein, exhibited an expanded distribution beyond canonical OPL boundaries in iPSC and W18 iKO organoids (Figure S5A). However, abundance metrics remained unchanged across all groups (Figures S5B–D).

Collectively, these data demonstrate that early *GRM6* deletion disrupts laminar consolidation and synaptic stratification of the OPL without reducing presynaptic ribbon abundance. Loss of mGluR6 prior to or during synaptogenesis leads to expansion or partial duplication of the synaptic layer, whereas deletion after synaptic maturation preserves OPL architecture.

### Early *GRM6* deletion selectively alters trans-synaptic scaffold assembly

We next asked whether disruption of OPL laminar architecture following *GRM6* deletion extends to trans-synaptic adhesion complexes that couple photoreceptor terminals to ON-bipolar dendrites. We focused on LRIT3 and ELFN2, two essential organizers of the photoreceptor–ON bipolar synapse.

At W32, LRIT3 immunoreactivity in control organoids formed a discrete punctate band within the OPL (Figure 4A). In both iPSC and W18 iKO organoids, LRIT3 puncta were spatially dispersed across the broadened or partially stratified synaptic zone, paralleling the laminar disorganization observed for presynaptic ribbon markers. In contrast, W26 iKO organoids exhibited a control-like LRIT3 distribution, with puncta largely restricted to the OPL. Quantification revealed no significant differences in LRIT3 puncta density across groups (control: 143.6 ± 13.5; iPSC iKO: 138.9 ± 13.3; W18 iKO: 139.4 ± 13.5; W26 iKO: 137.6 ± 15.6 puncta per 10□ µm²) (Figure 4B), indicating that overall LRIT3 abundance is preserved despite its spatial redistribution in early knockout conditions.

**Figure 4.**
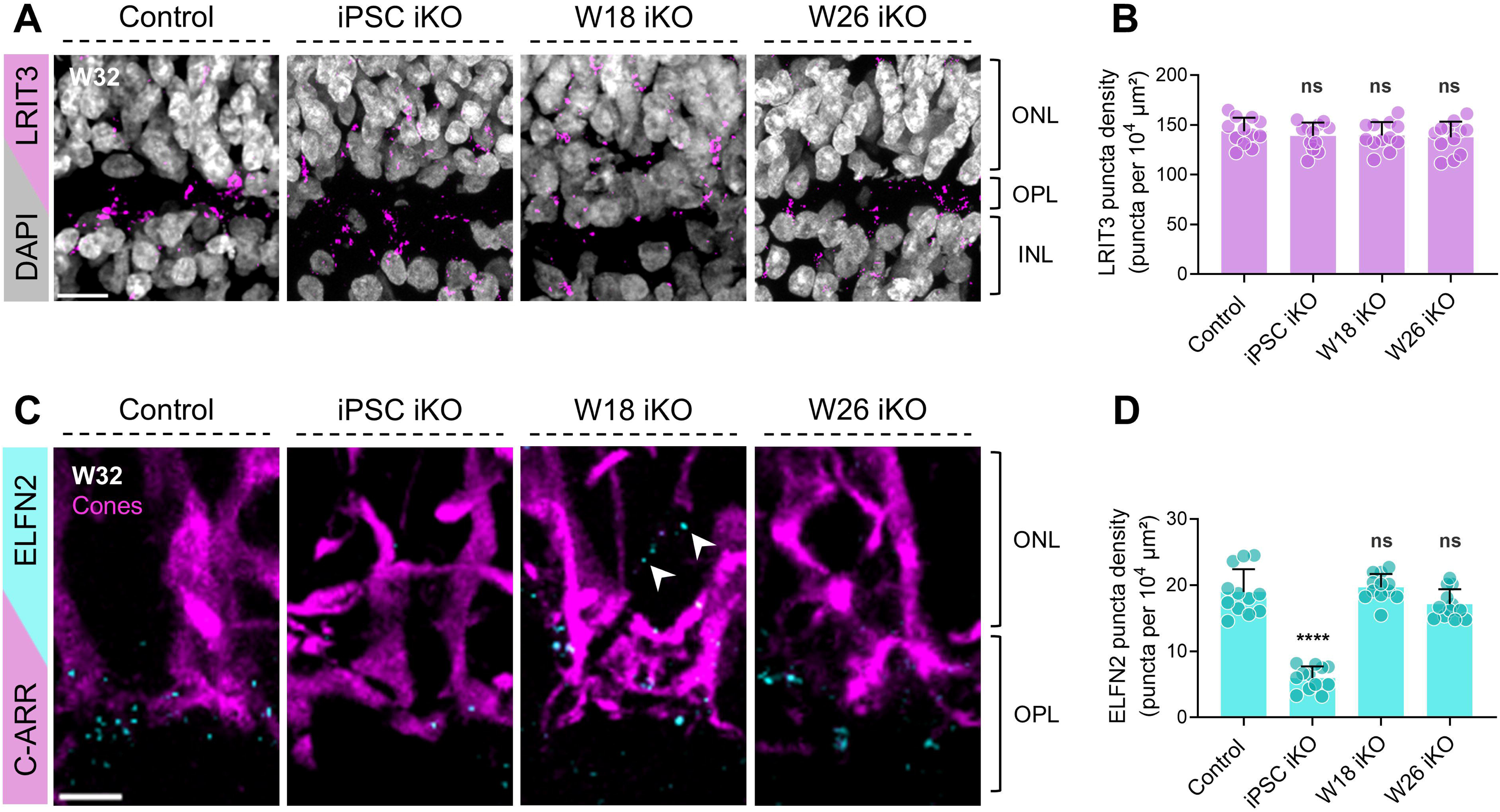
Early *GRM6* deletion selectively alters trans-synaptic scaffold assembly. (A) Immunofluorescence analysis of LRIT3 (magenta) in week 32 (W32) inducible knockout (iKO) human retinal organoids (HROs) following stage-specific *GRM6* deletion. ONL, outer nuclear layer; OPL, outer plexiform layer; INL, inner nuclear layer. Nuclei are counterstained with DAPI (gray). Scale bar, 10 µm. (B) Quantification of LRIT3 protein puncta density within standardized regions of interest (ROIs) spanning ONL, OPL, and adjacent INL. (C) Immunofluorescence analysis of ELFN2 (cyan) co-stained with a cone marker (cone-arrestin, C-ARR; magenta) in W32 iKO HROs following stage-specific *GRM6* deletion. White arrowheads indicate mislocalized ELFN2 within the pseudo-OPL. Nuclei are counterstained with DAPI (gray). Scale bar, 5 µm. (D) Quantification of ELFN2 protein puncta density within standardized ROIs spanning ONL, OPL, and adjacent INL. Quantifications were performed on 12 organoids per group from 3 independent differentiations. Four ROIs per organoid were analyzed, and values represent the mean ± SD per organoid (n = organoids). Statistical comparisons were performed using one-way ANOVA with Dunnett’s multiple-comparison test relative to the control. ****P < 0.0001.

We then examined ELFN2, a cone-associated trans-synaptic organizer enriched at cone terminals (Figure 4C). In control organoids, ELFN2 puncta were confined to the OPL and closely apposed to cone terminals. Strikingly, iPSC-stage *GRM6* deletion led to a marked reduction in ELFN2 signal, with sparse puncta detected within the synaptic layer. Quantitative analysis confirmed a significant decrease in ELFN2 puncta density in iPSC iKO organoids (5.9 ± 1.7 puncta per 10□ µm²) compared to controls (18.9 ± 3.4 puncta per 10□ µm²) (Figure 4D). By contrast, ELFN2 densities in W18 and W26 iKO organoids remained indistinguishable from controls (19.6 ± 2.0 and 17.1 ± 2.2 puncta per 10□ µm², respectively), although W18 samples displayed a broader spatial distribution consistent with the duplicated OPL architecture.

Collectively, these data reveal that early *GRM6* loss differentially compromises trans-synaptic scaffold components. Whereas LRIT3 abundance is preserved despite its spatial redistribution, ELFN2 requires early *GRM6* expression for its successful integration into the synaptic complex.

### Early *GRM6* deletion impairs the postsynaptic ON-bipolar signaling complex

Having established that early *GRM6* loss disrupts presynaptic architecture and trans-synaptic scaffolding, we next asked whether these structural defects compromise the assembly of the postsynaptic signaling machinery within ON-bipolar cell dendrites (Figure 5).

**Figure 5.**
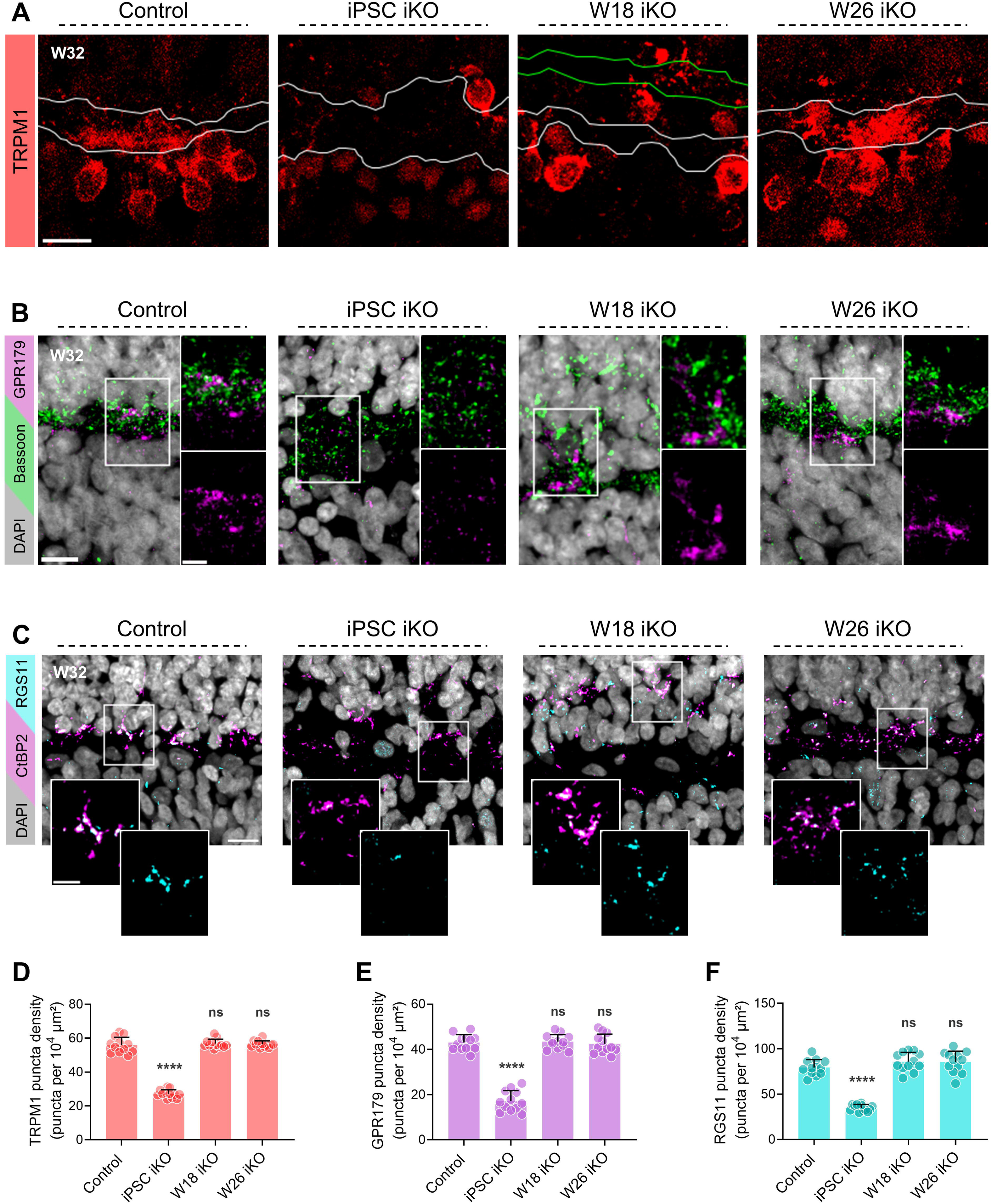
Early *GRM6* deletion impairs the postsynaptic ON-bipolar signaling complex. (A) Immunostaining of TRPM1 (red) in week 32 (W32) inducible knockout (iKO) human retinal organoids (HROs) following stage-specific *GRM6* deletion. The primary and pseudo OPL are indicated by gray and green lines, respectively. Scale bar, 10 μm. (B) Immunostaining of GPR179 (magenta) co-stained with presynaptic ribbon marker Bassoon (green) in W32 iKO HROs following stage-specific *GRM6* deletion. Nuclei were counterstained with DAPI (gray). Scale bar, 10 μm; Inset 5 μm. (C) Immunostaining of RGS11 (cyan) co-stained with presynaptic ribbon marker CtBP2 (magenta) in W32 iKO HROs following stage-specific *GRM6* deletion. Nuclei were counterstained with DAPI (gray). Scale bar, 10 μm; Inset 5 μm. (D-F) Quantification of postsynaptic protein puncta density within standardized ROIs spanning ONL, OPL, and adjacent INL (puncta per 10□ μm²): (D) TRPM1 puncta density, (E) GPR179 puncta density, and (F) RGS11 puncta density. Quantifications were performed on 12 organoids per group derived from three independent differentiations (n = organoids). Four ROIs per organoid were analyzed, and values represent the mean ± SD per organoid. Statistical comparisons were performed using one-way ANOVA with Dunnett’s multiple-comparison test relative to the control.****P < 0.0001.

We first analyzed TRPM1, the cation channel that mediates ON-bipolar depolarization downstream of mGluR6 (Figure 5A). In control W32 organoids, TRPM1 was enriched along the plasma membrane of ON-bipolar cell bodies and dendritic shafts, with strong accumulation at dendritic tips within the OPL. A similar membrane-associated and OPL-enriched distribution was preserved in W26 iKO organoids. In contrast, iPSC-stage *GRM6* deletion profoundly altered TRPM1 localization. OPL-associated signal was markedly reduced, plasma membrane enrichment was strongly diminished, and TRPM1 appeared diffusely distributed within the cytoplasm of ON-BCs. Quantification confirmed a significant reduction in TRPM1 puncta density in iPSC iKO organoids (27.2 ± 2.3 puncta per 10□ µm²) compared to controls (55.9 ± 4.6) (Figure 5D). Deletion at W18 produced a distinct phenotype; membrane localization and OPL enrichment were largely preserved, but TRPM1-positive dendritic tips were redistributed across both the primary OPL and the pseudo-OPL stratum. Despite this spatial duplication, overall TRPM1 puncta density in W18 iKO organoids (56.8 ± 2.5 puncta per 10□ µm²) was indistinguishable from controls.

We next examined GPR179, a scaffold protein required for anchoring the mGluR6 signaling complex at ON-BC dendritic tips (Figure 5B). In control organoids, GPR179 puncta were sharply confined to the OPL and co-localized with presynaptic Bassoon. Similar to TRPM1, GPR179 was significantly reduced in iPSC iKO organoids (17.1 ± 4.5 versus 43.0 ± 3.5 puncta per 10□ µm² in controls), whereas W18 and W26 iKO samples exhibited densities comparable to the control group (Figure 5E). However, W18 iKO organoids displayed spatial redistribution consistent with the duplicated OPL architecture.

Finally, we assessed the localization of RGS11, a regulator of Gα_o_ signaling that accelerates response termination at ON-bipolar dendritic tips. RGS11 was co-labeled with the presynaptic ribbon marker CtBP2 to evaluate its alignment with photoreceptor synaptic sites (Figure 5C). In control organoids, RGS11 puncta were tightly confined to the OPL and closely apposed to CtBP2-positive ribbons. This pattern was maintained in W26 iKO organoids. In contrast, iPSC-stage deletion resulted in a marked reduction of RGS11 within the OPL and loss of its tight alignment with CtBP2-positive ribbons.

RGS11 puncta density was significantly decreased (34.9 ± 3.7 puncta per 10□ µm²) compared to controls (79.3 ± 8.7) (Figure 5F). In W18 iKO organoids, total RGS11 abundance (85.8 ± 10.2 puncta per 10□ µm²) remained comparable to controls; however, RGS11 puncta were redistributed across both the primary and pseudo-OPL strata. Together, these findings reveal a stage-dependent requirement for *GRM6* in the assembly and structural integrity of the postsynaptic ON-bipolar signaling complex. Deletion at the pluripotent stage results in loss and mislocalization of core signaling components, and deletion during early synaptogenesis (W18) produces a related phenotype characterized by preserved protein abundance but clear laminar mislocalization. In contrast, deletion after synaptogenesis preserves both postsynaptic protein abundance and localization.

### Cone axon mistargeting underlies OPL disorganization following early *GRM6* deletion

Given that early *GRM6* deletion disrupts synaptic lamination and alters the ON-bipolar signaling complex, we next asked whether structural defects in photoreceptor axon targeting underlie the broadened and duplicated OPL observed in early knockout organoids.

In control organoids, cone arrestin (C-ARR)–positive cones extended axons basally and terminated within a defined region of the OPL adjacent to the INL, consistent with the normal axon termination patterns, ^36^ where they aligned with GNG13-positive ON-bipolar dendritic tips (Figure 6A). In contrast, iPSC-stage *GRM6* deletion caused cone axon mistargeting. Many cone axons failed to reach the INL-adjacent region of the OPL and instead terminated prematurely within more apical regions of the synaptic zone. ON-bipolar dendrites extended toward these ectopic cone terminals, producing the observed expansion of the OPL. Deletion at W18 produced an intermediate phenotype. In duplicated OPL regions, cone targeting defects were partial rather than global; while some cones projected correctly to the primary OPL, others terminated within an upper pseudo-layer, contributing to duplication of the synaptic band. ON-bipolar dendrites extended toward both strata, consistent with the duplicated ribbon and postsynaptic distributions described above. In contrast, W26 iKO organoids exhibited largely preserved cone axon targeting and maintained a consolidated OPL comparable to controls.

**Figure 6.**
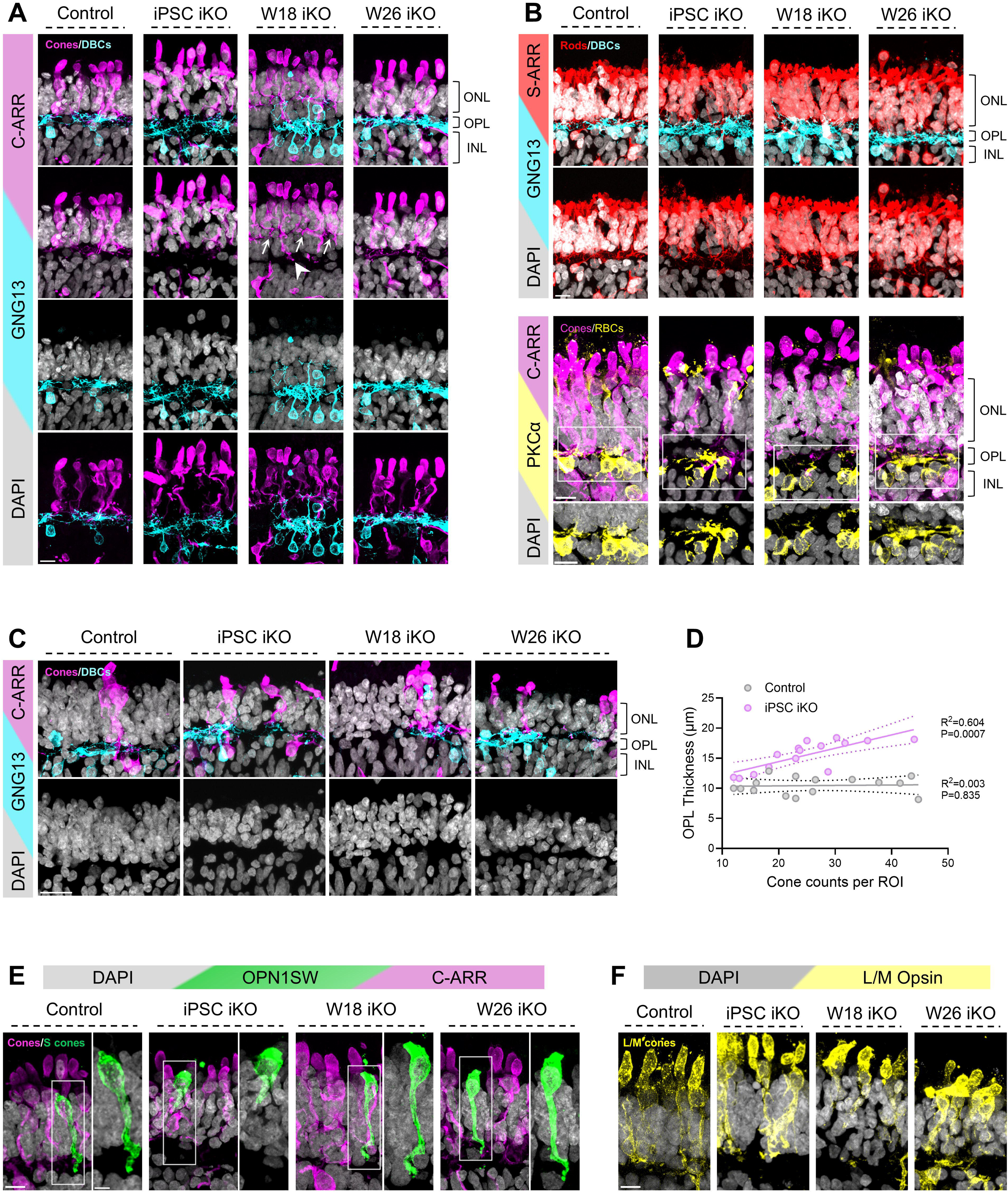
Cone axon mistargeting underlies OPL disorganization following early *GRM6* deletion. **(A)** Representative confocal images of week 32 (W32) inducible knockout (iKO) human retinal organoids (HROs) stained for cones (C-ARR; magenta), and ON-bipolars (GNG13; cyan) following stage-specific *GRM6* deletion. Arrowheads indicate correctly targeted cone terminals, whereas arrows indicate mistargeted cone terminals within the pseudo-OPL in the W18 iKO group. Nuclei were counterstained with DAPI (gray). Scale bar, 10 μm. **(B)** Representative confocal images of W32 iKO HROs stained for rods (S-ARR; red), ON-bipolars (GNG13; cyan), and rod bipolars (PKCα; yellow) following stage-specific *GRM6* deletion. Nuclei were counterstained with DAPI (gray). Scale bar, 10 μm. **(C)** Representative confocal images of W32 cone-poor iKO HROs stained for cones (C-ARR; magenta), and ON-bipolars (GNG13; cyan) following stage-specific *GRM6* deletion. Nuclei were counterstained with DAPI (gray). Scale bar, 20 μm. **(D)** Scatter plot showing the relationship between cone number per ROI and OPL thickness measured in control and iPSC iKO HROs. Five organoids per group were selected, spanning three cone-density categories (10–20 cones per ROI, 20–30 cones per ROI, and >30 cones per ROI). **(E)** Representative images of W32 iKO HROs stained for S cones (OPN1SW; green) and cones (C-ARR; magenta) following stage-specific *GRM6* deletion. Nuclei were counterstained with DAPI (gray). Scale bar, 10 μm; Inset 5 μm. **(F)** Representative images of W32 iKO HROs stained for L/M cones (L/M opsin; yellow) following stage-specific *GRM6* deletion. Nuclei were counterstained with DAPI (gray). Scale bar, 10 μm.

To determine whether this was specific to the cone photoreceptors, we examined rod circuitry. S-arrestin–positive rod axons extended normally into the OPL across all genotypes, and PKCα-positive rod bipolar dendrites remained appropriately confined to the original OPL without aberrant elongation (Figure 6B).

If cone mistargeting drives OPL expansion and duplication, phenotypic severity may correlate with cone abundance, which is inherently variable across HROs. Consistent with this prediction, cone-poor iPSC and W18 iKO organoids exhibited attenuated laminar disorganization compared to cone-rich counterparts (Figures 6A and 6C). Quantitative analysis revealed a strong positive correlation between cone number per ROI and OPL thickness in iPSC iKO organoids (R² = 0.604, P = 0.0007), whereas no such relationship was observed in controls (R² = 0.003, P = 0.835) (Figure 6D). Regression slopes differed significantly between groups (F = 12.64, P = 0.0015), supporting a cone-dependent phenotype in which ectopic cone terminals actively expand the synaptic layer.

We next asked whether specific cone subtypes were responsible for the OPL widening and splitting. In iPSC-stage iKO organoids, both S cones (OPN1SW-positive) and L/M cones displayed targeting defects. S cones exhibited severe axonal shortening and failed to project into the OPL (Figure 6E), whereas L/M cones terminated within more apical regions of the synaptic zone rather than the canonical INL-adjacent region of the OPL (Figure 6F). In contrast, W18 iKO organoids exhibited subtype-specific effects. S cones projected normally and reached the appropriate synaptic layer with morphology comparable to controls (Figure 6E). Mistargeting was restricted to a subset of L/M cones, whose axons terminated within an ectopic synaptic layer and contributed to the formation of a duplicated OPL (Figure 6F). W26 iKO organoids exhibited largely preserved subtype-specific cone morphology.

Collectively, these findings support a model in which cone axon mistargeting is a primary structural contributor to OPL expansion and duplication following early *GRM6* deletion.

### Early *GRM6* deletion disrupts inner retinal lamination associated with cone axon mistargeting

To determine whether cone axon mistargeting affects other retinal cell types, we examined the positioning of rods and inner retinal neurons in W32 organoids across all knockout conditions (Figure 7). A schematic overview illustrates the expected laminar positioning of rod, horizontal cell, and bipolar cell nuclei within the retina relative to the OPL under control conditions (Figure 7A).

**Figure 7.**
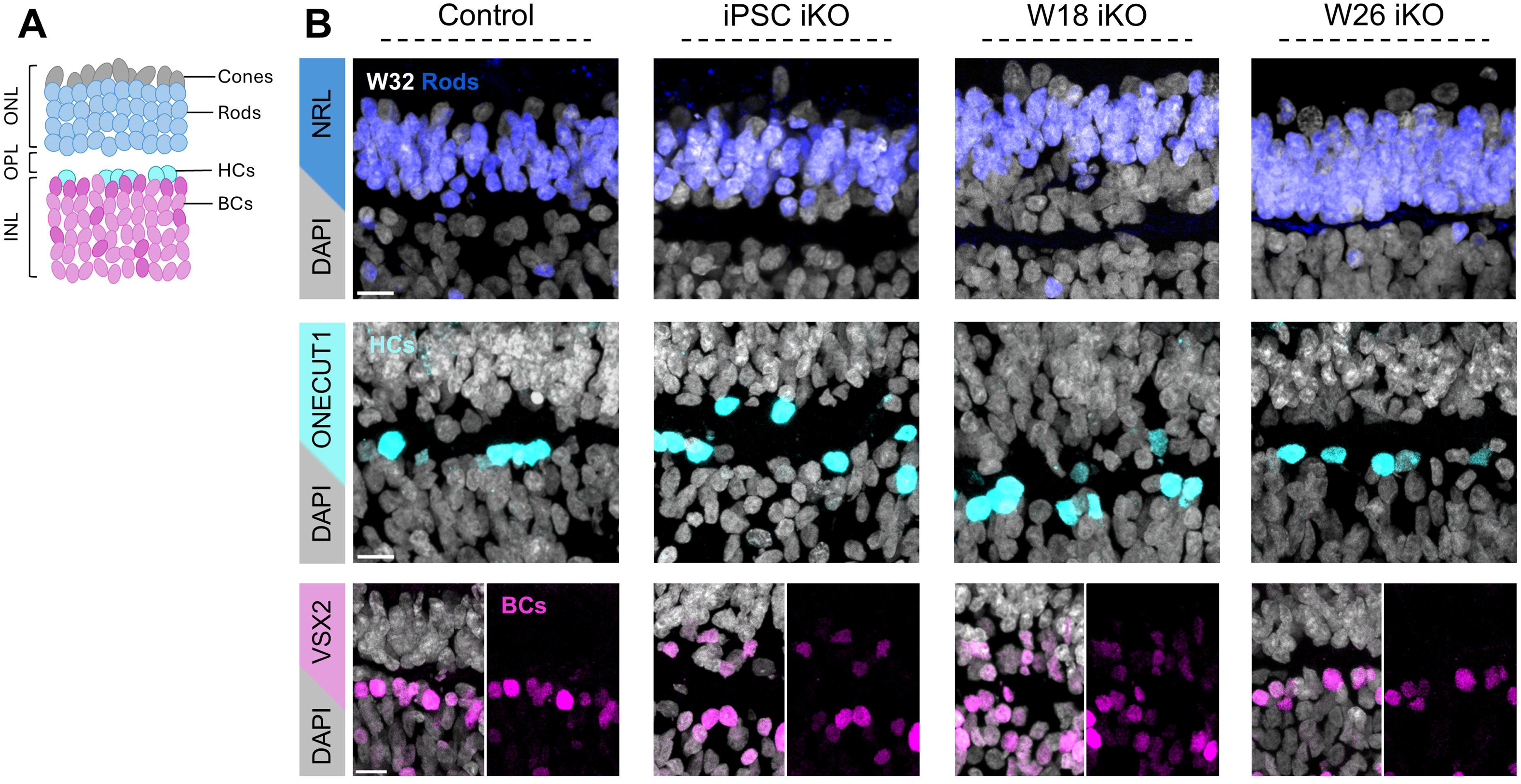
Early *GRM6* deletion disrupts inner retinal lamination associated with cone axon mistargeting. (A) Schematic illustrating the expected laminar positioning of rod photoreceptor, horizontal cells, and bipolar cell nuclei relative to the outer plexiform layer (OPL) under control conditions. (B) Representative immunofluorescence images of week 32 (W32) inducible knockout (iKO) human retinal organoids (HROs) following stage-specific *GRM6* deletion, stained for rod photoreceptors (NRL, blue), horizontal cells (ONECUT1, cyan), and bipolar cells (VSX2, magenta). Nuclei are counterstained with DAPI (gray). Scale bar, 10 μm. See also Figure S6

Immunostaining for NRL revealed that rods remained correctly localized within the ONL in all groups, showing no evidence of migration into the OPL or INL (Fig. 7B). ONECUT1-positive horizontal cells formed a discrete layer at the OPL interface in control and W26 iKO organoids but appeared more spatially dispersed in iPSC and W18 iKO groups, suggesting a broader laminar distribution under early knockout conditions. VSX2-positive bipolar cells exhibited more pronounced alterations. In iPSC and W18 iKO organoids, bipolar cells were less tightly confined to the INL and extended beyond their typical laminar boundaries (Figure 7B). Notably, there was a greater accumulation of VSX2-positive cells within ectopic synaptic regions in W18 iKO organoids, consistent with the more structured laminar duplication observed at this stage. This mislocalization is consistent with the redistribution of TRPM1-positive ON-bipolar cells observed in Figure 5A, where the TRPM1 signal extends into the ONL. In addition, immunostaining for GRIK1 revealed mislocalization of OFF-bipolar cells in iPSC and W18 iKO organoids (Figure S6), indicating that both ON and OFF bipolar populations are affected. By comparison, W26 iKO organoids retained a compact bipolar cell layer similar to controls.

Together, these findings indicate that early *GRM6* deletion disrupts inner retinal lamination in association with cone axon mistargeting, while rod lamination remains preserved.

## DISCUSSION

Our study reveals a previously unrecognized, temporally restricted role for mGluR6 in coordinating cone axon targeting and synaptic layer assembly during human retinal development. Although mGluR6 is classically defined as a postsynaptic receptor mediating ON-bipolar signaling,^3,25,26^ we show that it is transiently expressed in human cone photoreceptors and is required for proper cone axon targeting and laminar organization of the OPL. Using temporally controlled genetic ablation in human retinal organoids, we found that *GRM6* deletion prior to synaptogenesis leads to cone-dependent expansion or partial duplication of the OPL, whereas deletion after synaptic maturation preserves laminar architecture. These findings expand the role of mGluR6 from a simple signal transducer to a stage-specific regulator of synaptic assembly.

Studies in *Grm6* knockout mice report largely preserved retinal lamination, with phenotypes restricted to the loss of ON-bipolar physiological signaling.^10,37^ In the rod-dominant mouse retina, *Grm6* expression is confined to ON-bipolar dendrites, and no presynaptic role in photoreceptors has been described. Consistent with this, the defects in postsynaptic protein expression and localization observed in our early knockout conditions are likely explained by the canonical postsynaptic expression of mGluR6 in ON-bipolar cells and align well with phenotypes reported in mouse models.^14,38–40^ In contrast, our human organoid system also exhibits cone axon mistargeting, mislocalization of presynaptic proteins, and eventual OPL disorganization, features not reported in murine models. This divergence is explained by a key species-specific feature: the transient expression of mGluR6 in human cone photoreceptors during mid-gestation, a pattern that is faithfully recapitulated in human retinal organoids (Figure 1). Together, these findings highlight a developmental role for mGluR6 that is not predicted from rod-dominant mouse models and underscore the value of human systems for uncovering species-specific mechanisms of retinal circuit assembly.

The transient expression of mGluR6 in cones suggests a presynaptic developmental role distinct from its canonical postsynaptic function. While mGluR6 is often considered retina-specific, transcripts have been identified in non-neuronal tissues, including the kidney, corneal endothelium, and B lymphocytes,^41^ where it likely functions in non-synaptic capacities such as regulating calcium influx or cell proliferation.^42,43^ In our system, however, mGluR6 appears to serve a dual role as both a morphogenic scaffold and a glutamate-sensitive autoreceptor. In the broader CNS, Group III mGluRs (mGluR4, 7, and 8) typically function as presynaptic autoreceptors that provide feedback inhibition of glutamate release by modulating Ca^2+^ channel activity and vesicle trafficking.^44–47^ In other systems, mGluR signaling, specifically mGluR1, is known to regulate cytoskeletal dynamics and growth cone motility by modulating Rho GTPase activity.^48,49^ This model is consistent with our observation that early *GRM6* deletion leads to cone axon shortening and premature termination in apical regions of the OPL. The loss of mGluR6-mediated sensing or physical anchoring appears to prevent cone axons from reaching their appropriate basal targets. We found that early *GRM6* deletion disrupts the expression and localization of ELFN2, a known trans-synaptic binding partner.^50^ While ELFNs are typically characterized as agents that recruit mGluR6 to the postsynaptic site,^50–52^ our data suggest a bidirectional dependency in humans, suggesting that presynaptic mGluR6 acts as a prerequisite anchor for stabilizing the trans-synaptic scaffold. Without this positioning signal, the cone-bipolar junction fails to stabilize at the correct depth, resulting in the observed duplication or expansion of the synaptic layer.

Our data suggest that cone photoreceptors are the primary cellular drivers of the observed OPL defects. Cone axon mistargeting closely parallels the onset of laminar disorganization, whereas rod circuitry remains structurally intact across all conditions (Figure 6). This cone-dependent mechanism is further supported by our observation that lamination defects are significantly attenuated in cone-poor organoids. whereas bipolar cells exhibit pronounced mislocalization and horizontal cells show subtle dispersion, these alterations are best interpreted as secondary to the displacement of cone terminals. This model, in which mispositioned photoreceptor terminals drive broader laminar instability, is supported by knockouts of specific non-canonical Wnt signaling components in the mouse retina, where the mistargeting of rod terminals alone is sufficient to drive broader laminar instability and generate a duplicated OPL.^53^ Although the rod pathway remains stable in our human system, the shared phenotype suggests a general rule: the precise positioning of photoreceptor terminals provides the scaffold for outer retinal lamination.

The structural phenotypes observed in our study, particularly bipolar dendrite extension and synaptic disorganization, are consistent with a potential presynaptic contribution of mGluR6 in cone photoreceptors. Similar dendritic extension phenotypes have been reported in mouse models affecting presynaptic or structural components of the photoreceptor synapse. For example, mutations in the presynaptic calcium channel Cav1.4 (CACNA1F) or disruption of Spectrin-mediated cytoskeletal scaffolding lead to abnormal bipolar dendrite extension and synaptic disorganization.^54,55^ However, in these models, dendritic sprouting is thought to reflect attempts to track or compensate for destabilized or retracting photoreceptor terminals. In contrast, in our system, cone terminals appear to remain positioned within both the primary and ectopic synaptic layers, and bipolar dendrites extend toward these stable but mislocalized targets. Thus, while the phenotypes share common features, our findings suggest a distinct mechanism in which altered photoreceptor terminal positioning, rather than terminal loss or retraction, drives dendritic remodeling, a result that supports a presynaptic role for mGluR6 in the establishment of cone terminal lamination.

The use of an inducible knockout strategy enabled us to decouple the developmental and maintenance roles of mGluR6. The preservation of laminar architecture following deletion at W26 indicates that mGluR6 is an essential architect during initial circuit assembly but becomes dispensable once synaptic organization is achieved. Within this developmental window, the specific nature of the OPL defect (expansion or duplication) is dictated by the timing of mGluR6 loss. Early deletion at the iPSC stage likely disrupts the very first stages of synaptic organization, including initial interactions between mGluR6 and its pre- and postsynaptic partners.^56^ Without this molecular tethering, nascent cone terminals and bipolar dendrites fail to coalesce, resulting in a diffuse, broadened synaptic zone. In contrast, deletion during active lamination (W18) permits the initial formation of a primary layer but triggers the formation of an ectopic, apical synaptic layer as later-maturing cones fail to target correctly.^57,58^ The temporal hierarchy of cone subtype development further refines this stage-dependent framework.^59,60^ S-cones, which differentiate early, exhibit the most severe axonal shortening following iPSC-stage deletion. Interestingly, L/M cones appear less severely affected in the iPSC-stage knockout than in the W18 deletion. In the former, they often reach the OPL despite mistargeting, whereas in the latter, they frequently terminate prematurely in apical regions. This suggests that the prolonged absence of mGluR6 from the start of development may allow for compensatory remodeling or partial adaptation that is not possible when the protein is abruptly removed during the peak of laminar consolidation. These results highlight that the final OPL architecture is a product of both the specific timing of molecular loss and the varying developmental trajectories of human cone subtypes.

Mutations in *GRM6* cause congenital stationary night blindness (CSNB), a condition traditionally attributed to impaired ON-bipolar signaling.^8,61^ Our findings raise the possibility that *GRM6* mutations also influence early developmental organization of human cone synapses. The absence of overt OPL abnormalities in patients likely reflects two factors. First, the micron-scale structural perturbations observed here may fall below the resolution of clinical imaging modalities like optical coherence tomography (OCT). This is consistent with *LRIT3* mutations, another cause of CSNB, where patients typically present with normal OCT scans despite histological evidence of dendritic extension and synaptic disorganization in animal models.^62–64^ Second, retinal development in vivo occurs within a complex, activity-dependent environment in which compensatory remodeling and homeostatic mechanisms may partially restore laminar organization despite early defects.^65,66^

In summary, our study identifies a stage-specific role for mGluR6 in organizing synaptic layer assembly in the human retina and supports a cone-dependent mechanism underlying this process. These findings suggest that transient, cell-type-specific expression of a canonical signaling receptor can shape neural circuit architecture and highlight the critical role of developmental timing in defining gene function within complex neural tissues.

### Limitations of the study

Although human retinal organoids recapitulate key aspects of retinal development, functional synaptic transmission was not directly assessed, and electrophysiological studies will be needed in the future to determine how cone mistargeting impacts circuit function. In addition, while our data support a cone-driven mechanism, definitive validation will require cone-specific *GRM6* deletion. Finally, the downstream mechanisms of presynaptic mGluR6 remain to be defined, including whether its developmental role is mediated through canonical signaling or scaffolding interactions.

## RESOURCE AVAILABILITY

### Lead contact

Further information and requests for resources and reagents should be directed to and will be fulfilled by the lead contact, Aaron Nagiel

### Materials Availability

Materials generated in this study are available upon request.

### Data and code availability

All data reported in this study will be shared by the lead contact upon request.

Any additional information required to reanalyze the data reported in this work is available from the lead contact upon request.

## ACKNOWLEDGEMENT

We thank the Children’s Hospital Los Angeles Stem Cell Analytics Core and Cellular Imaging Core for providing instrumentation and technical support. We are grateful to Kirill Martemyanov and David Cobrinik for providing the mGluR6 antibody and GNAT2 reporter organoids, respectively. We thank Rosanna Caldron for technical assistance and Mark Reid for advice on statistical analysis. Schematic illustrations were created using BioRender. This work was supported in part by an unrestricted grant to the Department of Ophthalmology at the Keck School of Medicine of USC from Research to Prevent Blindness (New York, NY, USA) (A.N.), a National Eye Institute Career Development Award (K08EY030924) (A.N.), the Las Madrinas Endowment in Experimental Therapeutics for Ophthalmology (A.N.), the Knights Templar Eye Foundation (A.N.), the Donald E. and Delia B. Baxter Foundation (A.N.), and a Research to Prevent Blindness Career Development Award (A.N.). L.B. was supported by the California Institute for Regenerative Medicine (CIRM) training grant (EDUC4-12802) through the CHLA Training Program for Stem Cell and Regenerative Medicine Research.

## AUTHOR CONTRIBUTIONS

Conceptualization, A.N. and L.B.; methodology, L.B., P.G., H.H., J.B., S.P.B., M.C., K.S., M.E.T., and B.T.G.; investigation, L.B., A.N., and H.H.; funding acquisition, A.N. and L.B.; supervision, A.N.; writing – original draft, L.B.; writing – review and editing, A.N and LB with input from all authors.

## DECLARATION OF GENERATIVE AI AND AI-ASSISTED TECHNOLOGIES

The authors used generative AI tools to assist with language editing and text refinement. The authors reviewed and edited the output and take full responsibility for the content of the publication.

## SUPPLEMENTAL INFORMATION

Document S1. Figures S1–S6 and Tables S1 and S2.

## STAR METHODS

### KEY RESOURCES TABLE

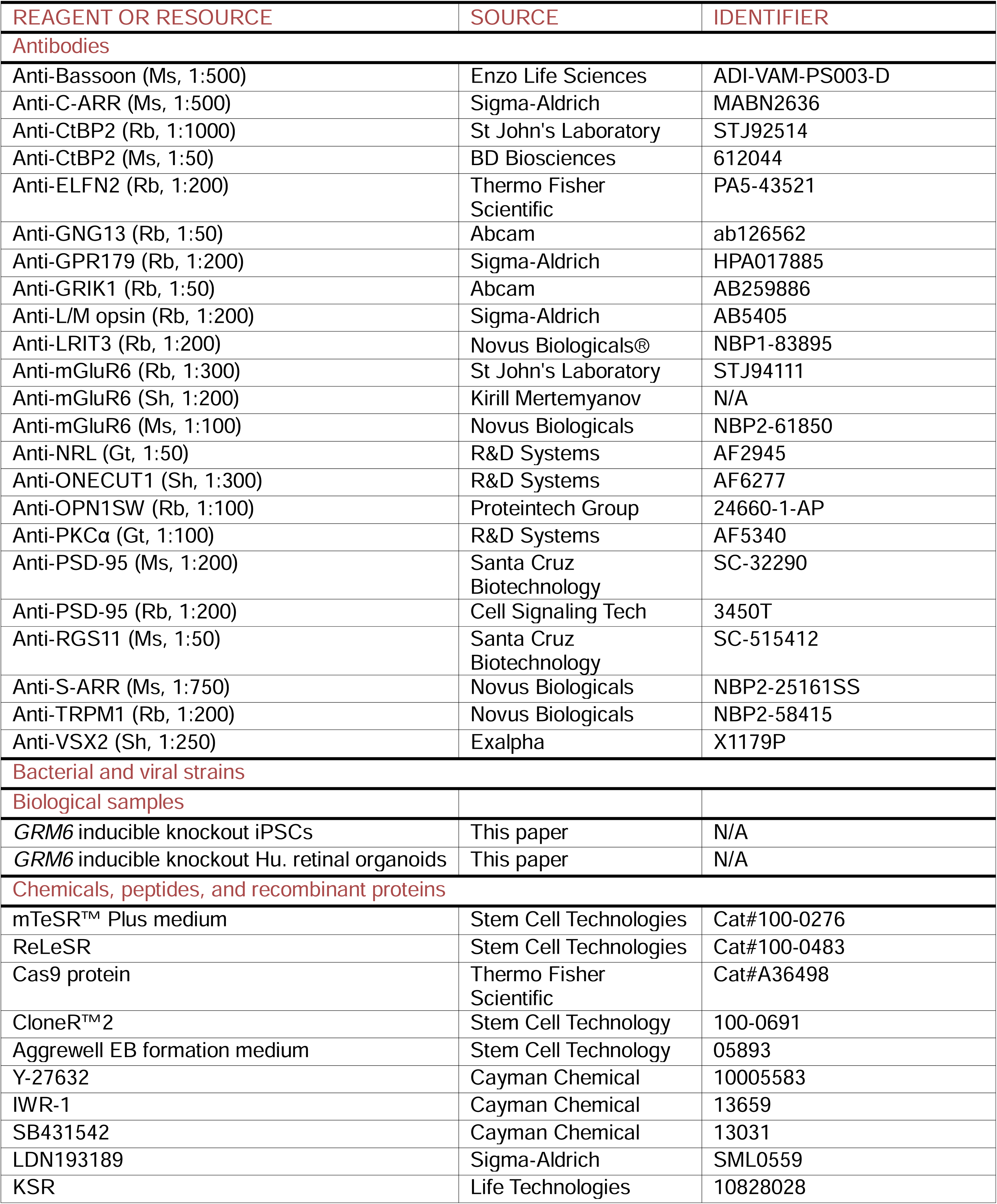
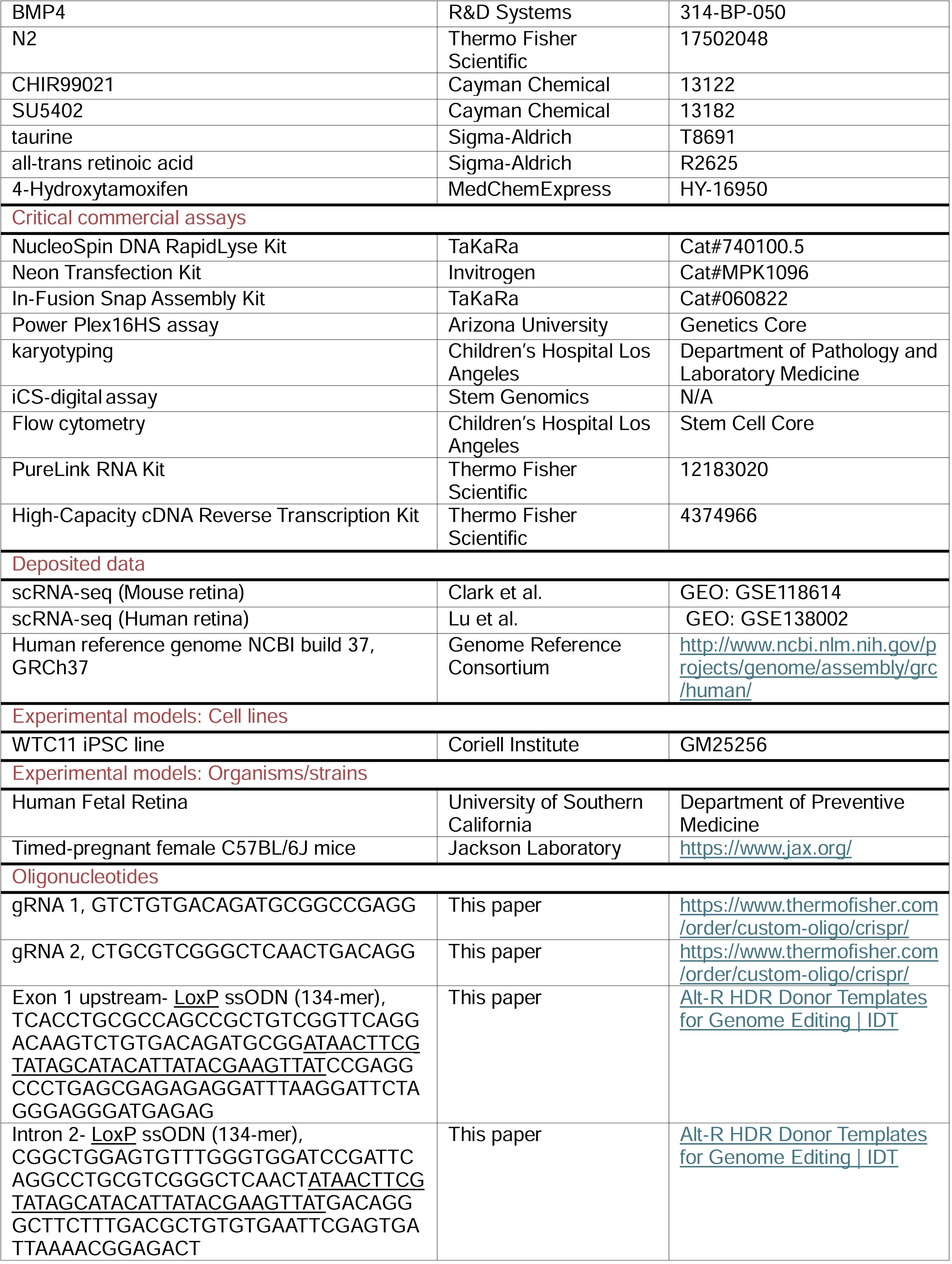
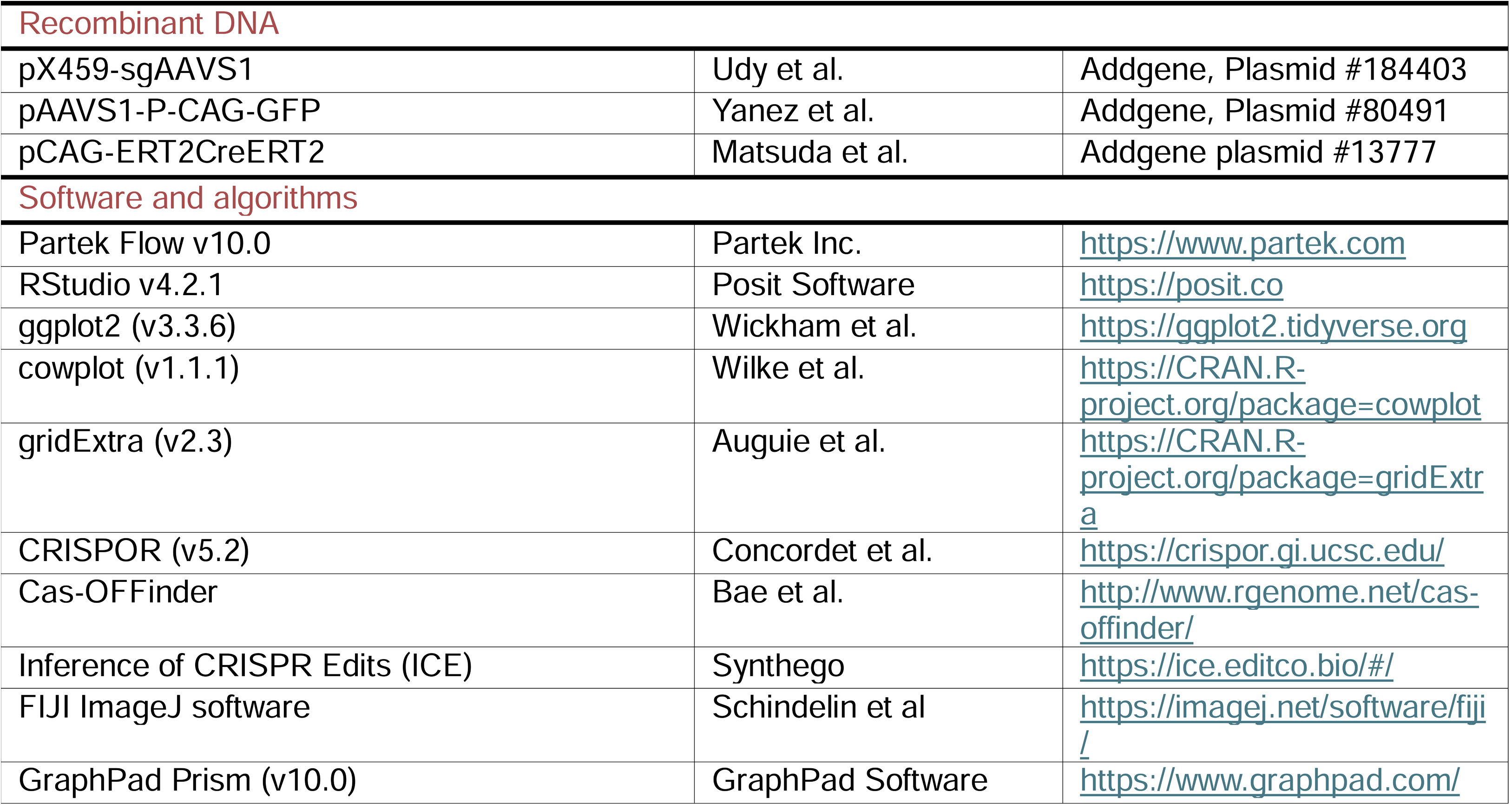

### EXPERIMENTAL MODEL AND STUDY PARTICIPANT DETAILS

#### Human fetal retinal tissue

Human fetal retinal tissues were obtained from elective terminations of pregnancy conducted with full informed consent. Tissue procurement was performed under protocols approved by the Institutional Review Boards of the University of Southern California (USC-HS-13-0399) and Children’s Hospital Los Angeles (CHLA-14-2211, CHLA-15-00363). Donors were informed about the purpose of the research, the specific procedures that would be conducted, and the potential future use of the donated tissues in research. Participation was entirely voluntary and did not influence the clinical decision regarding termination. Only gestational age and the presence or absence of genetic or structural abnormalities were collected to protect donor anonymity. This research falls under Category 1b of the ISSCR 2021 Guidelines for Stem Cell Research and Clinical Translation. All procedures were conducted following a suitable research oversight process and in accordance with the principles of the Declaration of Helsinki. No gametes, embryos, or embryo models were created or used in this study.

#### Mouse retinal tissue

All animal experiments were conducted under research protocol #79-17, approved by the Animal Research Committee at Children’s Hospital Los Angeles, and in strict accordance with the NIH Guide for the Care and Use of Laboratory Animals. Timed-pregnant C57BL/6J mice were obtained from The Jackson Laboratory. Eyes were collected from pups between P6 and P12, as well as from adult mice.

#### Human iPSC line

The iPSC line WTC11 (Coriell Institute, GM25256) was used as the parental line for generating a *GRM6* inducible knockout (iKO) line via CRISPR-Cas9 gene editing. This study complies with the 2021 ISSCR guidelines. iPSC work was conducted under the oversight of the Stem Cell Research Oversight Committee (SCRO ID: STUDY00002969) at Children’s Hospital Los Angeles, and informed consent for the original derivation of the WTC11 line is on file through the supplying institution.

### METHOD DETAILS

#### Single-cell RNA-seq data analysis

Single-cell RNA sequencing (scRNA-seq) count matrices and corresponding cell annotations from publicly available datasets of mouse and human retina were obtained from the Gene Expression Omnibus (GEO) repository.^23,24^ Data processing and analysis were performed using Partek Flow v10.0.^67^ Genes expressed in fewer than 0.01% of total cells within each dataset were excluded to minimize noise from lowly expressed transcripts. Normalization was conducted by scaling raw counts to counts per million (CPM), followed by log_₂_ transformation with an offset of 1 to stabilize variance. For visualization, bubble plots were generated in RStudio v4.2.1 employing the ggplot2 (v3.3.6), cowplot (v1.1.1), and gridExtra (v2.3) packages.^68–71^ In these plots, bubble color intensity corresponds to the mean normalized expression level of each gene, while bubble size indicates the proportion of cells expressing that gene. Both expression and percentage scales were independently normalized within each dataset to facilitate direct comparison.

#### Half-life prediction

The mGlR6 potential *in vivo* half-life was estimated in *Homo sapiens* and *Mus musculus* using computational prediction and experimental proxy approaches. Computational estimates were generated using a protein language model–based regression framework (PLTNUM) with both sequence-based (ESM2) and structure-aware (SaProt) encoders, as well as an empirical N-end rule–derived approximation implemented in ProtParam.^31,32^ Human and mouse mGluR6 sequences were obtained from UniProt (O15303 and Q5NCH9).^72^ As experimental proxies, published half-life measurements for related group III metabotropic glutamate receptors were compiled from dynamic SILAC and SILAM-based proteome turnover datasets (PRIDE PXD008514 and TissuePPT).^33,73^ Brain-region values were averaged to generate a single half-life estimate per protein, and the median half-life was calculated across all prediction outputs and proxy measurements to provide a consensus estimate

#### Guide RNA design

Guide RNAs (gRNAs) targeting the 5′ region of *GRM6* exon 1 (∼100 bp upstream) and intron 2 (∼100 bp downstream of exon 2), referred to as gRNA1 and gRNA2, respectively, were designed using the CRISPOR online tool (v5.2).^29^ The genomic sequence spanning the target region was retrieved from the human reference genome NCBI build 37, GRCh37,^74^ and uploaded to CRISPOR to identify candidate gRNAs. The top five-ranked gRNAs were further evaluated using Cas-OFFinder to assess potential off-target sites, focusing on mismatch positions and the likelihood of DNA:RNA loop formation. gRNAs with the fewest predicted off-target effects, particularly those with fewer than three mismatches, were selected for subsequent genome editing experiments.^75^

#### Single-stranded oligodeoxynucleotide design

Single-stranded oligodeoxynucleotides (ssODNs) were designed to serve as donor templates for the insertion of loxP sites using CRISPR/Cas9-mediated genome editing. Approximately 500 bp of genomic sequence flanking each Cas9 cut site (corresponding to each gRNA target region) was retrieved from the GRCh37 human genome assembly via NCBI^74^ and sequenced to confirm the absence of single-nucleotide polymorphisms (SNPs) that could interfere with Cas9 activity. Primer sequences are listed in Table S1. The 34 bp loxP sequence was positioned directly at the Cas9 cleavage site, flanked by 50 bp homology arms on either side, resulting in a total ssODN length of 134 bp.

#### Human pluripotent stem cell culture

The WTC11 iPSC line (Coriell Institute, GM25256) was maintained under feeder-free conditions in mTeSR Plus medium (Stem Cell Technologies, 100-0276) on Matrigel (Corning, 354277)-coated dishes. Cells were seeded at a density of 120,000 cells per 35 mm dish, fed every other day, and passaged every 5 days at approximately 80% confluency. For passaging, iPSC colonies were washed with Dulbecco’s Phosphate-Buffered Saline (DPBS; Corning, 21-031-CV) and gently dissociated by a 2-minute incubation with ReLeSR (Stem Cell Technologies, 100-0483) at 37°C. Following neutralization with mTeSR Plus medium, cells were centrifuged at 300 × g for 4 minutes and subsequently reseeded at the desired densities.

#### Biallelic knock-in of loxP sites

For targeted insertion of loxP sites, approximately 100,000 Accutase (Life Technologies, A1110501)-dissociated iPSCs were seeded per well in a 24-well plate and electroporated using the Neon Transfection System (Invitrogen, MPK5000) with pre-assembled ribonucleoprotein (RNP) complexes. Each RNP complex contained 12 pmol Cas9 protein (Thermo Fisher Scientific, A36498) and 12.5 pmol of gRNA, co-delivered with 18 pmol of the corresponding loxP-containing ssODN. Genomic DNA was isolated three days post-electroporation using the NucleoSpin DNA RapidLyse Kit (TaKaRa, 740100.5), and the targeted locus was amplified via PCR (PrimeSTAR GXL Premix, TaKaRa, R051A). Amplicons were subjected to Sanger sequencing using the forward primers listed in Table S1. Editing efficiency was assessed in bulk populations using the Inference of CRISPR Edits (ICE) analysis tool,^76^ confirming successful loxP integration. Single-cell clones were then isolated by limiting dilution, and clonal lines were expanded over 10 days. Biallelic insertion was confirmed by Sanger sequencing and ICE analysis, demonstrating 100% knock-in efficiency. The entire workflow was repeated for the insertion of the second loxP site, ensuring it was integrated in the same orientation as the first.

#### Monoallelic knock-in of ERT2-Cre-ERT2

A guide RNA and Cas9-expressing plasmid targeting the AAVS1 locus was obtained from Addgene (Plasmid #184403). The donor plasmid was constructed by replacing GFP in the pAAVS1-P-CAG-GFP backbone (Addgene plasmid #80491) with ERT2-Cre-ERT2 cDNA, which was amplified from pCAG-ERT2-Cre-ERT2 (Addgene plasmid #13777), using In-Fusion Snap Assembly (TaKaRa, 060822). iPSCs harboring floxed exons 1 and 2 of the *GRM6* gene were co-transfected with 2.2 µg of the donor plasmid and 1.7 µg of the AAVS1 gRNA/Cas9-expressing plasmid. Electroporated cells were cultured in mTeSR Plus medium supplemented with CloneR2 (Stem Cell Technology, 100-0691) for 72 hours, followed by selection with 350 ng/mL puromycin (Sigma, P7255) for 36 hours. After a 10-day recovery period, single clones were screened by PCR for monoallelic integration of ERT2-Cre-ERT2 into the AAVS1 locus. Genotyping employed two primer pairs: pair 101/102 flanking the AAVS1 left homology arm, yielding a 1.4 kb product for the unedited allele, and pair 103/104 including the puromycin resistance gene, yielding a 1.6 kb product for correctly targeted knock-in clones (See Table S1 for primer details).

#### Post-editing quality controls in edited iPSCs

Potential off-target sites for each guide RNA were identified using the Cas-OFFinder prediction tool. Genomic regions corresponding to the top five predicted off-target sites for each gRNA were amplified by PCR in established edited, hereafter inducible knockout (iKO), iPSCs and subjected to Sanger sequencing to assess potential off-target cleavage. Detailed information regarding the selected off-target sites is provided in Table S2. Authentication, genomic stability, and pluripotency of iKO iPSCs were assessed using the PowerPlex 16 HS Assay, G-banding karyotyping, and iCS-digital assay, and flow cytometry, respectively. Genomic stability was evaluated through both chromosomal analysis and targeted digital assays. Pluripotency was confirmed by flow cytometry using OCT3/4 as an internal nuclear marker and SSEA-4 as an external surface marker. All assays were performed by the respective service providers using their validated protocols.

#### Differentiation of human retinal organoids

Human retinal organoids (HROs) were generated from iKO iPSCs using a protocol adapted from our previously published method.^19^ On day 0 (D0), iPSCs were dissociated using Accutase and seeded into round-bottom 96-well plates at a density of 24,000 cells per well in 150 μL of AggreWell medium (Stem Cell Technologies, 05893), supplemented with 20 μM Y-27632 (Cayman Chemical, 10005583) and an ISL cocktail to promote embryoid body formation. The ISL cocktail contained 3 μM IWR-1 (Cayman Chemical, 13659), 10 μM SB431542 (Cayman Chemical, 13031), and 0.1 μM LDN193189 (Sigma-Aldrich, SML0559). From D1 to D5, cultures were maintained in gfCDM medium composed of 45% Ham’s F12 (Thermo Fisher Scientific, 11765047), 45% IMDM (Thermo Fisher Scientific, 12440046), 10% Knockout Serum Replacement (KSR; Life Technologies, 10828028), 1× Chemically Defined Lipid Concentrate (Gibco, 11905-031), 0.5× GlutaMAX (Life Technologies, 35050061), 450 μM monothioglycerol (Sigma-Aldrich, M6145), and 1× Penicillin-Streptomycin (Corning, 30-002-CI), supplemented with the ISL cocktail. On D6, 1.5 nM BMP4 (R&D Systems, 314-BP-050) was added to promote neural induction. Partial medium changes with fresh gfCDM were performed on D9, D12 (50%), and D15 (75%). From D19 to D23, cultures were transitioned to RPE induction medium containing DMEM/F12 (Thermo Fisher Scientific, 21331020), 1× N2 supplement (Thermo Fisher Scientific, 17502048), 1× GlutaMAX, 1× Penicillin-Streptomycin, 3 μM CHIR99021 (Cayman Chemical, 13122), and 5 μM SU5402 (Cayman Chemical, 13182). Beginning on D23, organoids were cultured in RDM3S-KZ medium [DMEM/F12 supplemented with 10% FBS, 1× N2, 1× GlutaMAX, 1× Penicillin-Streptomycin, and 0.5× Fungizone], with 0.1 mM taurine (Sigma-Aldrich, T8691) added starting on D30. On the same day, organoids were transferred to HEMA-coated 48-well plates to promote adhesion and maturation. From D37 to D42, the medium was gradually transitioned to a maintenance medium (MM) adapted from Zhong et al. Specifically, cultures received a 2:1 ratio of RDM3S-KZ to MM on D37, 1:2 on D40, and were fully transitioned to 100% MM on D42. MM consisted of a 1:1 mixture of DMEM (VWR, 54000-305) and DMEM/F12, supplemented with 1× B27 (Thermo Fisher Scientific, 12587010), 1× non-essential amino acids (NEAA), 1× GlutaMAX, 1× Penicillin-Streptomycin, 1× Fungizone (Omega Scientific, FG-70), 10% FBS, and 0.1 mM taurine. From D72 to D100, MM was supplemented with 1 μM all-trans retinoic acid (Sigma-Aldrich, R2625), which was reduced to 0.5 μM after D100 to support further photoreceptor maturation. Organoids were maintained in this medium throughout the culture. *GNAT2*-reporter human retinal organoids were generously provided by David Cobrinik (Children’s Hospital Los Angeles). These organoids were generated as previously described.

#### Inducible knockout strategy and validation

Three inducible knockout treatment paradigms were implemented to disrupt *GRM6* expression. In Group 1, edited iPSCs were treated with 1□μM 4-hydroxytamoxifen (4-OHT; MedChemExpress, HY-16950) for two consecutive days before the initiation of retinal organoid differentiation. The 4-OHT-containing medium was refreshed daily, and treatment was discontinued at the onset of differentiation. The resulting organoids are hereafter referred to as iPSC iKO HROs. In Group 2, retinal organoids derived from iKO iPSCs were treated at differentiation week 18 (W18) with 4 μM 4-OHT for seven consecutive days, with daily media replacement; these are referred to as W18 iKO HROs. Group 3 received an identical treatment regimen at differentiation week 26 and is referred to as W26 iKO HROs. Each experimental group was accompanied by a matched vehicle-treated control group to account for non-specific effects of the treatment. To evaluate *GRM6* inducible knockout efficiency at the transcript level, total RNA was extracted from control and iKO HROs using the PureLink RNA Kit (Thermo Fisher Scientific, 12183020), and cDNA was synthesized using the High-Capacity cDNA Reverse Transcription Kit (Thermo Fisher Scientific, 4374966). Quantitative PCR (qPCR) was performed using SYBR Green chemistry on a QuantStudio 7 Flex Real-Time PCR System (Thermo Fisher Scientific) with 1□μL cDNA per reaction in triplicate. Gene expression was normalized to the geometric mean of the internal reference gene. See Table S1 for primer details

#### Tissue collection, embedding, and cryosectioning

Tissue collection, embedding, and sectioning of mouse and human retina were performed as described previously.^77^

Mouse retina: Eyes were collected from postnatal day (P) 10–12 pups. To maintain RNase-free conditions, all instruments were cleaned with ELIMINase (VWR, 10498-216), and buffers were prepared using DEPC-treated water. Eyes were rinsed three times in 1× PBS (Thermo Fisher Scientific, AM9625) and fixed in 4% paraformaldehyde (PFA; VWR, 100503-917; 1 mL per eye). The cornea was carefully removed under a dissecting microscope, and the lens was extracted using forceps. Adult eyes were post-fixed in 4% PFA for an additional 15 minutes, followed by PBS washes. Eyecups were cryoprotected in 15% sucrose overnight at 4□°C, then in 30% sucrose for 3□hours. Samples were embedded in O.C.T. compound (VWR, 25608-930) using 10 × 10 mm cryomolds, frozen on dry ice, and stored at −80□°C. 30 μm cryosections were cut in dorsoventral orientation and stored at −80□°C.

Human retina: Donor eye tissue was processed under RNase-free conditions. A 12-well plate was prepared with 70% ethanol (EtOH) and 1× PBS. Tissue was immersed in 70% EtOH for 5□seconds, then washed three times in PBS. In cold PBS, residual tissue was trimmed, and the cornea was punctured with a needle and removed using Vannas scissors. The lens was gently extracted, and the eyecup was fixed in 4% PFA for 1□hour at room temperature, followed by three 5-minute PBS washes. Cryoprotection and embedding were performed as described for mouse tissue. Serial dorsoventral sections (10□μm) were cut and stored at −80□°C.

Human retinal organoids (HROs): Control and tamoxifen-treated HROs were collected at W32 and cryoprotected in 30% sucrose overnight at 4□°C. Organoids were embedded in O.C.T. using 10 × 10 mm cryomolds, frozen on dry ice, and stored at −80□°C. To ensure matched processing, control and iKO HROs were cryosectioned simultaneously, with alternating sections (e.g., control in section 1 and tamoxifen-treated in section 2) mounted on the same slide. Serial sections (20□μm) were stored at −80□°C until use.

#### RNA fluorescence in situ hybridization (RNA-FISH)

Probe sets were custom-designed and synthesized by Molecular Instruments, Inc. using target mRNA accession numbers and organism information to ensure high hybridization efficiency and specificity. Multiplexed RNA-FISH was performed on sectioned mouse retina, human retina, and human retinal organoids following the manufacturer’s protocol with minor adaptations. Slides were equilibrated at room temperature (RT) for 10 minutes, fixed with 4% PFA, and washed three times in PBS. Sections were then placed in cold 70% ethanol in a Coplin jar and stored overnight at −20□°C. The following day, slides were air-dried for 5 minutes, outlined with a PAP pen, and washed twice with 2× saline sodium citrate (SSC; Invitrogen, 15557044). Prehybridization was carried out for 30 minutes at 42□°C in a humidified chamber. Probes (1.6 pmol per 100 μL hybridization buffer) were prepared at 37□°C, applied to sections, and incubated overnight in a 42□°C oven. Unbound probes were removed with four 5-minute washes in wash buffer at 37□°C, followed by two 5-minute washes in 2× SSC at RT. Sections were incubated with amplification buffer for 30 minutes at RT. Hairpins (6 pmol each of hairpin 1 and hairpin 2) were heat-denatured at 95□°C for 90□seconds, cooled to RT for 30 minutes, and then added to sections and incubated overnight in a humidified chamber at RT. Excess hairpins were washed off with five washes in 2× SSC, followed by a final wash with 2× SSC containing 5□μg/mL DAPI (Thermo Fisher Scientific, D1306). Slides were mounted with coverslips using ProLong Gold Antifade Mountant (Thermo Fisher Scientific, P36970) and sealed for imaging. All RNA-FISH experiments were independently repeated three times.

#### Immunofluorescence staining (IF)

Frozen sections were thawed at room temperature and subsequently permeabilized and blocked for 1□hour in a humidified chamber at RT using 1× PBS containing 3% horse serum and 0.3% Triton X-100. Sections were then incubated with primary antibodies diluted in blocking buffer for 2□hours at RT. Following three washes with 1× PBS, slides were incubated with Alexa Fluor–conjugated secondary antibodies diluted in blocking buffer for 1□hour at RT. To visualize cell nuclei, 5□μg/mL DAPI was added to 1× PBS during the final wash step. Immunostained tissue sections were mounted with coverslips using ProLong Gold Antifade Mountant media (Thermo Fisher Scientific, P36970) and sealed. A list of primary and secondary antibodies used is provided in the Key Resources Table. All IF experiments were independently repeated three times.

#### Confocal fluorescence microscopy

Confocal z-stacks were acquired using a STELLARIS 5 system mounted on a DMi8 inverted microscope with an HC PL APO CS2 63×/1.4 oil immersion lens (Leica Microsystems). Laser lines at 405, 488, 555, and 639 nm were used to excite DAPI and Alexa Fluor 488, 568, and 647, respectively. The pinhole was set to 1 Airy unit for all channels, and voxel dimensions were 0.25 × 0.25 × 0.4 μm³ for 63× images. Laser power and detector gain were adjusted to maximize dynamic range while avoiding pixel saturation.

To minimize sampling bias in the analysis of HROs, which exhibit radial and laminated architecture, images were acquired from four evenly spaced positions corresponding to the 12, 3, 6, and 9 o’clock orientations of each organoid cross-section. This imaging strategy was applied consistently across all experimental groups and replicates.

Post-acquisition processing, including brightness adjustment, cropping, channel overlay, and maximum intensity projection, was performed using FIJI (ImageJ, NIH). Identical imaging settings and post-processing parameters were applied for each marker across all experimental conditions.

### QUANTIFICATION AND STATISTICAL ANALYSIS

All quantification graphs and statistical analyses were performed using GraphPad Prism (v10.6.0).

#### mRNA level quantification

Relative *GRM6* mRNA levels were quantified by qPCR from HRO samples obtained from three independent differentiation batches (n = 3 biological replicates per condition). Data are presented as mean ± SD. Statistical significance was determined using one-way ANOVA followed by Dunnett’s multiple comparisons test.

#### RNA-FISH puncta quantification

RNA-FISH puncta were quantified using FIJI (ImageJ, NIH).^78^ Individual RNA molecules were identified as diffraction-limited puncta using the Find Maxima function, with identical detection and thresholding parameters applied across all samples. Puncta with an apparent diameter of approximately 0.5 µm were considered single transcripts, whereas larger puncta were quantified as multiple transcripts based on their relative size. To account for differences in tissue size, puncta were quantified within defined 250 µm² regions of interest (ROIs).

#### OPL architecture quantification

OPL architecture was quantified using line intensity profiling in FIJI (ImageJ, NIH).^78^ For each ROI, five vertical line scans were drawn perpendicular to the ONL–INL axis using the straight-line tool with a fixed line width of 25 µm to minimize local intensity variability. Line positions were placed at five evenly spaced locations within each ROI and maintained at corresponding relative positions across all ROIs and groups to ensure unbiased sampling.

To quantify total OPL thickness, line profiles were generated in the DAPI channel to delineate nuclear layers. The OPL was defined as the DAPI-negative region between the ONL and INL. OPL thickness was measured as the distance between the ONL and INL boundaries along each line scan. For each ROI, the mean OPL thickness was calculated from the five line-scans.

To assess synaptic layer organization, line profiles were generated from the Bassoon channel along the same line positions. Intensity-versus-distance plots were examined to identify synaptic peaks. An ROI was classified as exhibiting a single OPL when a single dominant Bassoon intensity peak was detected between the ONL and INL boundaries. An ROI was classified as exhibiting a double OPL when two spatially separated Bassoon peaks were observed within the OPL region, defined by: 1. Each peak exceeding background intensity by at least twofold, 2. Peak-to-peak separation greater than 5 µm, and 3. Reproducible detection in at least three of five line scans per ROI. The percentage of ROIs exhibiting double OPL organization was calculated for each HRO.

#### Depth distribution quantification

Spatial distribution of Bassoon puncta relative to the ONL-OPL boundary was quantified using FIJI (ImageJ, NIH).^78^ A freehand line was drawn along the ONL boundary for each ROI to account for local curvature. The boundary was converted to a binary mask and used to generate a Euclidean distance map (Process → Binary → Distance Map). This produced a pixel-wise map in which each pixel value corresponded to its perpendicular distance (µm) from the ONL boundary. Bassoon puncta were then identified using consistent thresholding parameters across all samples. Diffraction-limited puncta were detected using Analyze Particles within a predefined size range corresponding to synaptic puncta. Centroid coordinates were extracted for each punctum. For each punctum, the corresponding distance value was extracted from the distance map at its centroid coordinate. This analysis generated a list of distances from the ONL boundary for each ROI. For each ROI, two parameters were calculated: (1) the mean distance from the ONL boundary, representing the average distance of Bassoon puncta relative to the ONL–OPL boundary regardless of direction, and (2) the interquartile range (IQR) of the signed distances, reflecting the dispersion of puncta across the OPL while preserving directional information.

#### Synaptic puncta quantification

Puncta density, size, and area fraction of synaptic proteins were quantified using FIJI (ImageJ, NIH).^78^ Background subtraction was performed using a rolling-ball radius optimized for each marker and then applied consistently across all images for that marker. To capture both correctly localized and ectopically distributed puncta, quantification was performed within rectangular ROIs measuring 250 µm × 80 µm, spanning the ONL, OPL, and the apical portion of INL. This approach allowed unbiased detection of synaptic proteins within the canonical OPL as well as mislocalized puncta extending into adjacent layers. Puncta were identified using consistent thresholding followed by the Analyze Particles function. Size filters were set to include diffraction-limited puncta corresponding to synaptic structures while excluding background noise. The same threshold and size parameters were applied to all images within each experimental marker. For each ROI, the following measurements were extracted: 1. Puncta density: number of puncta per ROI area (puncta/µm²), 2. Mean puncta size: average area (µm²) of detected puncta, and 3- Area fraction: cumulative puncta area divided by total ROI area.

#### Correlation analysis

Correlation analysis was performed to assess the relationship between cone number per ROI and OPL thickness. A total of 15 organoids per group were selected, with five organoids representing each of three cone-density categories (10–20 cones per ROI, 20–30 cones per ROI, and >30 cones per ROI). Correlation was evaluated using Pearson’s correlation coefficient.

