## Supplemental File for "mGluR6 coordinates cone terminal targeting and synaptic layer assembly during human retinal development"

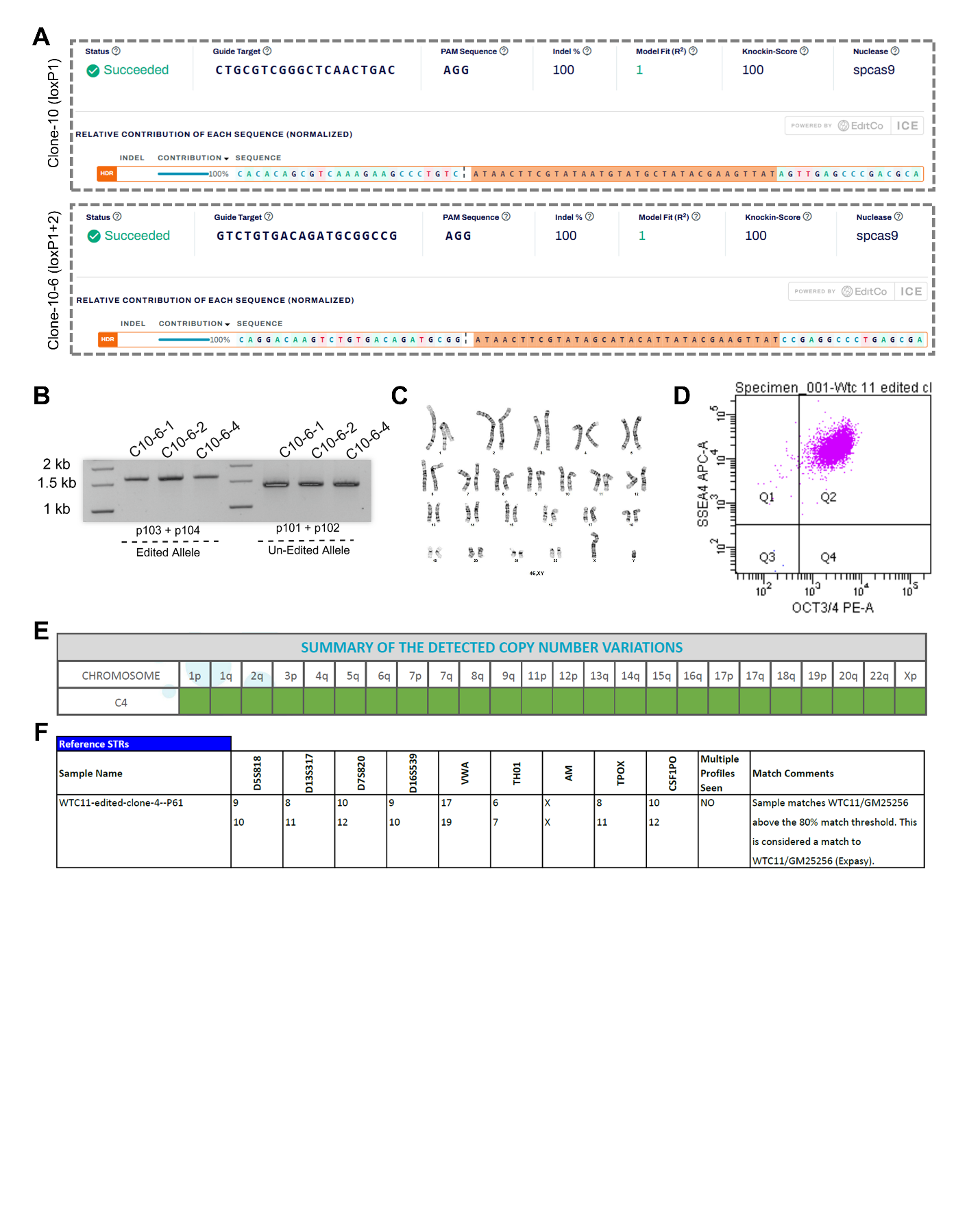

**Figure S1. Validation and quality control of a tamoxifen-inducible *GRM6* knockout human WTC-11 iPSC line. Related to Figure 2.** (A) Representative Sanger sequencing traces and Inference of CRISPR Edits (ICE) analysis confirming biallelic knock-in of loxP sites flanking *GRM6* exons 1 and 2 in single iPSC clones. (B) PCR genotyping strategy validating targeted integration of the ERT2-Cre-ERT2 cassette into the AAVS1 safe-harbor locus. Primer pair 101/102 amplified a 1.4 kb product from the un-edited allele, whereas primer pair 103/104 amplified a 1.6 kb product corresponding to the correctly targeted allele containing the puromycin resistance cassette. Three independent *GRM6*-iKO iPSC clones (10-6-1, 10-6-2, and 10-6-4) were identified; clone 10-6-4 (hereafter referred to as Clone-4) was selected for subsequent experiments. (C) G-banding karyotype analysis demonstrating normal chromosomal structure in Clone-4 *GRM6*-iKO iPSCs. (D) Flow cytometric analysis of pluripotency markers OCT3/4 and SSEA-4, indicating maintenance of an undifferentiated pluripotent state. (E) iCS-digital PCR assay confirming genomic stability and the absence of major copy number variations or aneuploidy. (F) Short tandem repeat (STR) profiling verifying the identity of the Clone-4 *GRM6*-iKO iPSC line using the PowerPlex 16 HS assay.

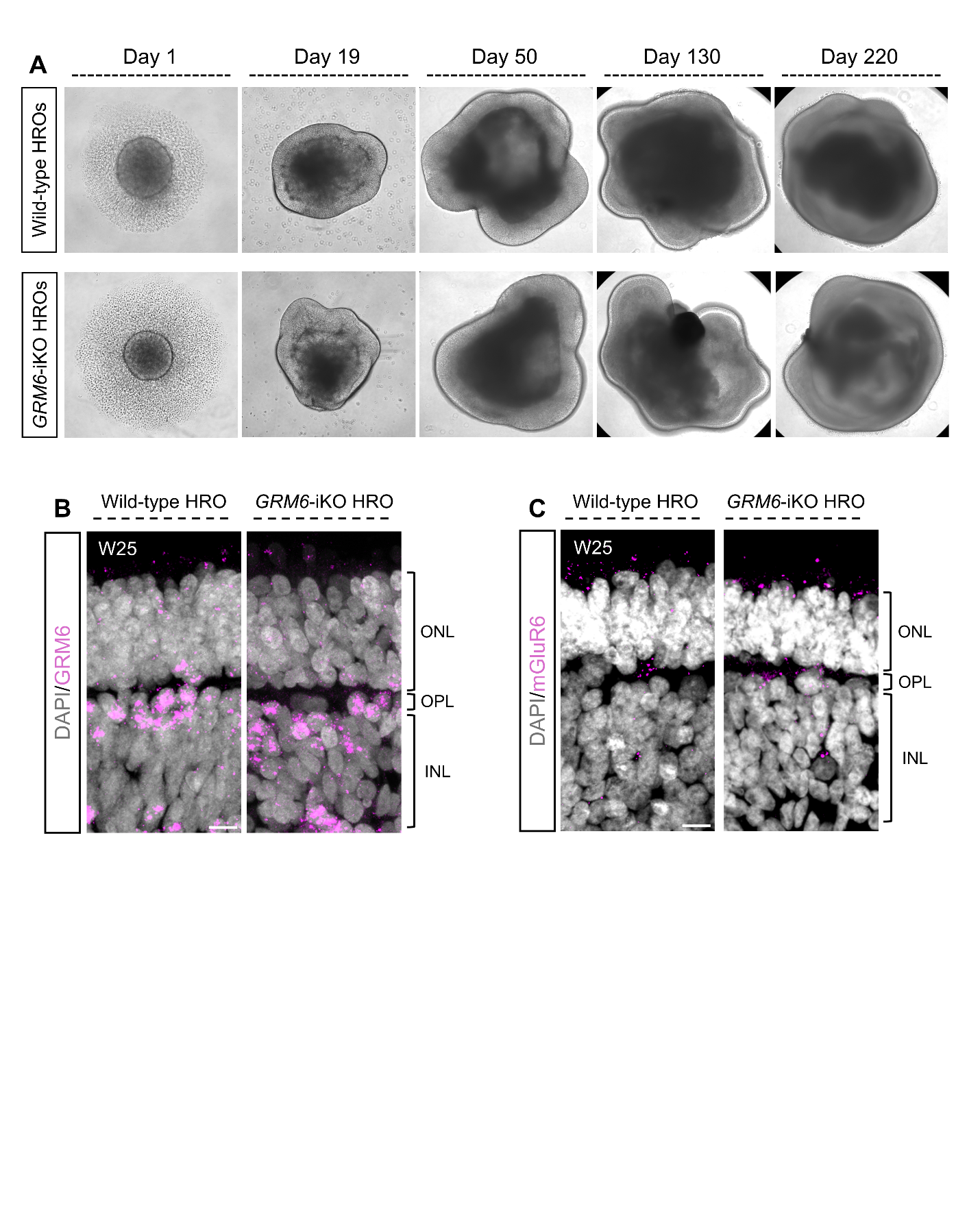

**Figure S2. *GRM6*-iKO iPSCs retain normal retinal differentiation capacity and baseline *GRM6* expression prior to Cre activation. Related to Figure 2.**

(A) Brightfield images showing morphological progression of wild-type and *GRM6*-iKO human retinal organoids (HROs) at representative stages of differentiation. (B) RNA-FISH for *GRM6* transcripts (magenta) in week 25 (W25) wild-type and *GRM6*-iKO HROs in the absence of tamoxifen induction. (C) Immunofluorescence staining for mGluR6 protein (magenta) in W25 wild-type and *GRM6*-iKO HROs without tamoxifen treatment. Nuclei are counterstained with DAPI (gray). Scale bars, 10 μm.

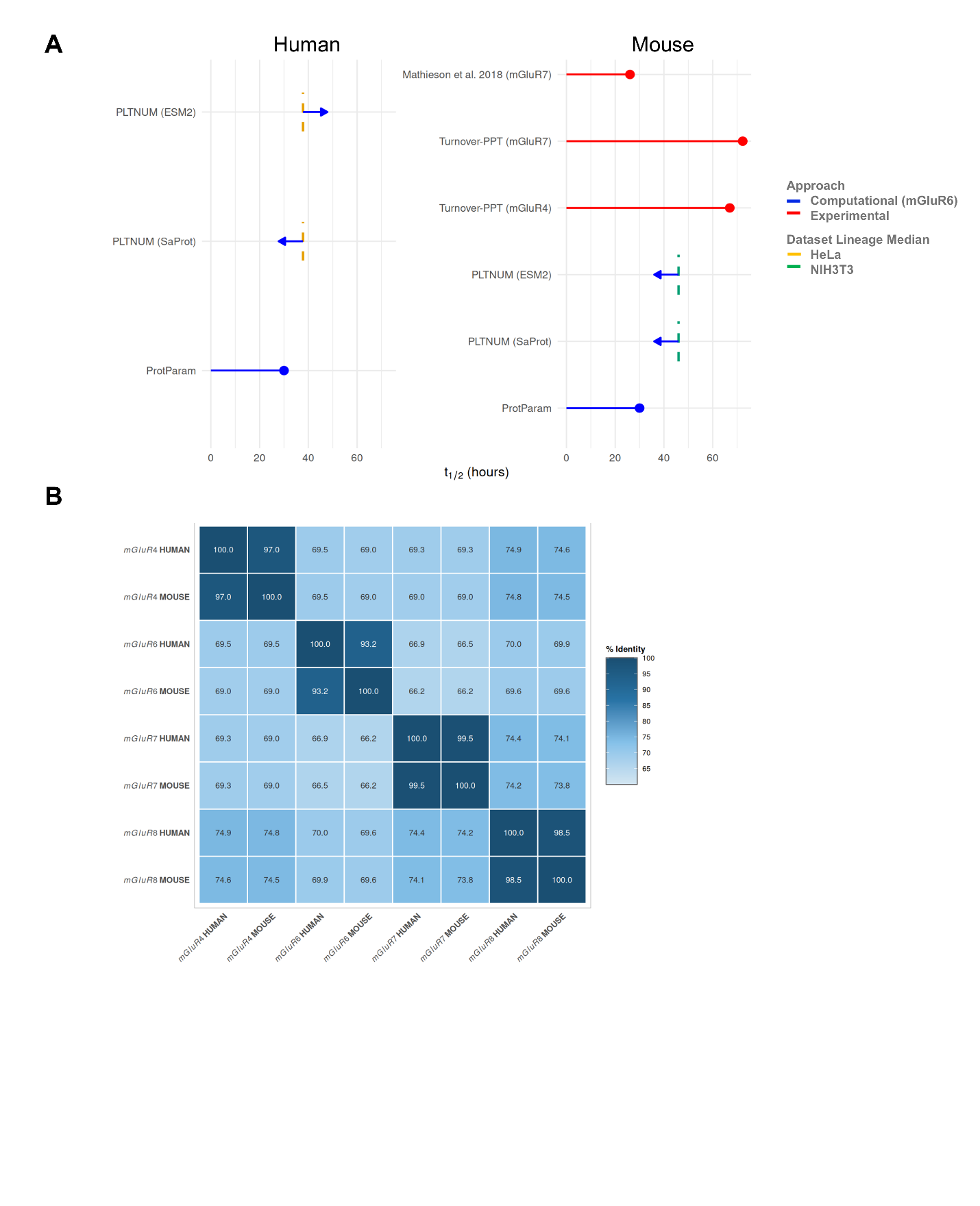

**Figure S3. mGluR6 half-life prediction and interspecies conservation. Related to Figure 2**.

(A) Estimated half-life (t₁/₂) of mGluR6 in human and mouse derived from computational prediction and experimental measurements of related Group III mGluRs. Computational predictions (blue) were generated using ProtParam and PLTNUM algorithms trained on independent proteomic turnover datasets. Arrows indicate predicted values with directional adjustment relative to dataset-specific lineage median half-lives (dashed vertical lines; HeLa and NIH3T3 reference datasets). Experimentally measured half-lives of related Group III mGluRs in mouse retina (red) are shown for comparison. (B) Percent identity matrix of human and mouse Group III mGluRs generated using Clustal Omega multiple sequence alignment. Heatmap values indicate pairwise amino acid sequence identity (%).

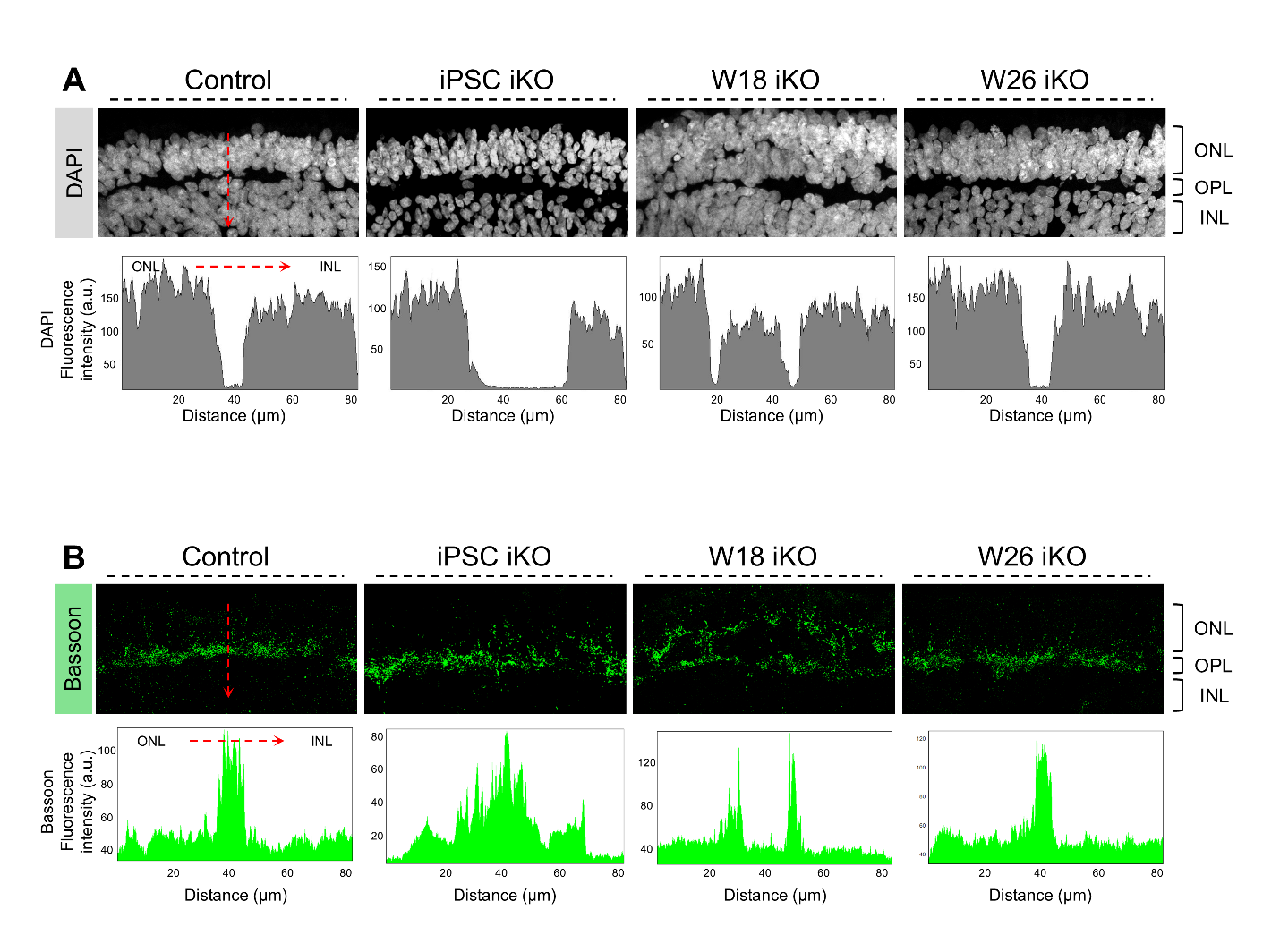

**Figure S4. Line profile analysis used to quantify OPL thickness and synaptic stratification. Related to Figure 3.**

(A) Representative example of vertical line (red dashed line) profiling across the ONL–OPL–INL axis. A series of standardized lines (25 µm width) were drawn perpendicular to the laminar axis at uniformly spaced intervals across each region of interest (ROI). DAPI intensity profiles were used to identify nuclear layer peaks corresponding to the ONL and INL, and total OPL thickness was defined as the distance between these peaks. (B) Representative Bassoon fluorescence intensity profiles from control, iPSC iKO, W18 iKO, and W26 iKO organoids. The number of dominant intensity peaks within each profile was used to classify ROIs as exhibiting a single OPL or a double (pseudo-duplicated) OPL.

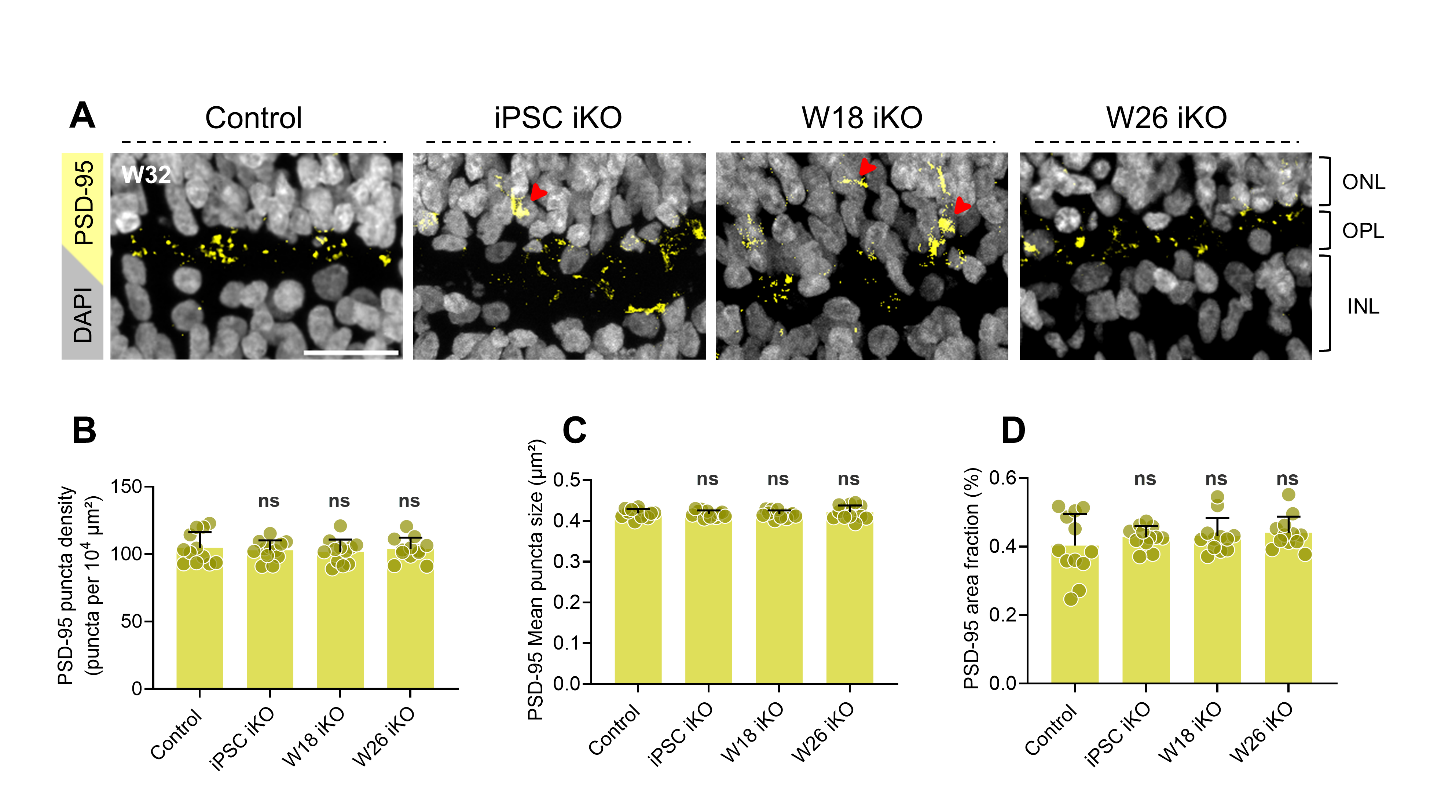

**Figure S5. Presynaptic protein PSD-95 distribution and quantification following stage-specific *GRM6* deletion. Related to Figure 3.**

(A) Representative immunofluorescence images of PSD-95 (yellow) in W32 HROs following *GRM6* deletion at the indicated developmental stages. Red arrowheads indicate mislocalized puncta in iPSC and W18 iKO HROs. Scale bar, 20 µm. (B–D) Quantification of PSD-95 puncta within standardized regions of interest (ROIs) spanning the ONL–OPL–INL. (B) Puncta density (puncta per 10⁴ µm²), (C) mean puncta size (µm²), and (D) area fraction (percentage of ROI area occupied by signal).

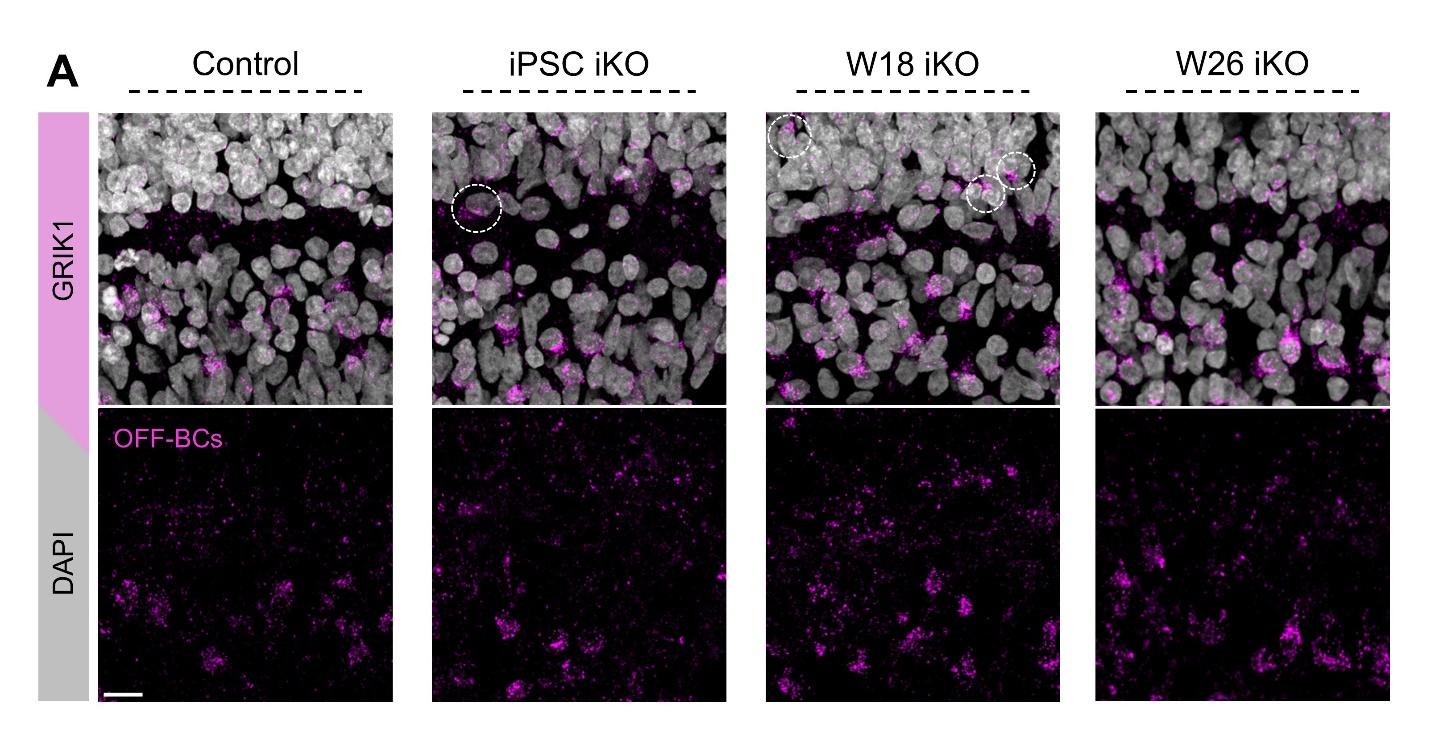

**Figure S6. Mislocalization of OFF-bipolar cells following early *GRM6* deletion. Related to Figure 7.**

Representative immunofluorescence images of iKO HROs at week 32 (W32) stained for the OFF-bipolar cell marker GRIK1 (magenta). Nuclei are counterstained with DAPI (gray). Mislocalized OFF-bipolar cells are indicated by dashed white circles. Lower panels show the corresponding GRIK1 signal alone. Scale bar, 10 μm.

**Table S1. Oligonucleotide sequences used in this study.**

| **Primers for the confirmation of SNP absence and LoxP site insertion** | |
| --- | --- |
| gRNA 1 primer forward | CTGCAAGGAGAAAGCTGGGG (used for sequencing) |
| gRNA 1 primer reverse | AGCGCCACGAGCAGC |
| gRNA 2 primer forward | TGTCCAGCGTCCCTCTCCC (used for sequencing) |
| gRNA 2 primer reverse | GAAGCCAGGCCTTCCTCCTT |
| **Primers for the confirmation of ERT2-Cre-ERT2 insertion** | |
| Primer 101 | TCGACTTCCCCTCTTCCGATG |
| Primer 102 | CTCAGGTTCTGGGAGAGGGTAG |
| Primer 103 | CATTGTCACTTTGCGCTGCC |
| Primer 104 | CGCTCGTAGAAGGGGAGGTT |
| **Primers for inducible knockout validation** | |
| 1.5 kb primer forward | GAGATTGGGAGAGGGAGGGG |
| 1.5 kb primer reverse | GCCCGGGTTCCAAACAAAAC |
| *GRM6* qPCR primer forward | TGCGCCTGTTTGCGATAC |
| *GRM6* qPCR primer reverse | GACACATAGTTCCATCCCAGTG |
| **Primers for amplifying gRNA1 off-target regions** | |
| Off-target 1 | F: TGCTGCAGTGCTATAACGAAAGA  R: GCCAATATACCCTGGTGCCTCT |
| Off-target 2 | F: GTCTAGTGAAGCTGGAGGGC  R: GGGGCTCAGTGTTTCTTCCA |
| Off-target 3 | F: GGCGGTGTCCACTTCTTCCT  R: GGGCCAGCTTAACCTTGGGA |
| Off-target 4 | F: AGTGGGGTCGGGAGAGACTT  R: CTGTCCCACCCCTGTTCTGG |
| Off-target 5 | F: GTTGGATTACAATCACAGGCTTATAC  R: CTGGTTCCTTCTCACTTGGT |
| **Primers for amplifying gRNA2 off-target regions** | |
| Off-target 1 | F: AGACTTCTTGGAAACAGTGGCA  R: CTGAGTGCCCTTCCCACAAC |
| Off-target 2 | F: AACGGGGAAGCTGTAGGGAG  R: CTTTGGGGCAGAGCTCAAGG |
| Off-target 3 | F: AAAGGTGGTCAGTGGAAGCC  R: CCACTCTCCAGGGCCTCAAA |
| Off-target 4 | F: TGGACTGGCACAGGAGAGAG  R: GAGAGGGGACAGACACCCTG |
| Off-target 5 | F: TGTTCTGCAACGTTCAGGTG  R: TCAATTCTTCTCCTCAGGATGTTCA |
| **Primers for amplifying AAVS1 gRNA off-target regions** | |
| Off-target 1 | F: CTGGCTGTGACCACTCACCA  R: GTCGACCCCTCGGGCTAATC |
| Off-target 2 | F: ATCATGGTGAGGGGACAGCG  R: TGGGTTGCTTTCCCAGTGGT |
| Off-target 3 | F: GGGTCTCTGCTCTGGAAACC  R: CTCGTGCAAGTTGCGTACAG |
| Off-target 4 | F: TCAGTGAAGGAAGCCATGTGGT  R: GGCCCCCATGTCTGTACGAT |
| Off-target 5 | F: GCTCCCAAGAGCCAACTCCA  R: TAGCACCGGGCAAGCATTTG |

**Table S2. Top five off-targets predicted by Cas-OFFinder CRISPR-Cas9 gRNA checker**

| **gRNA1 Off-targets** | |
| --- | --- |
| Off-target 1 | CTCTGTGACAGCATGCTGCCTCGG, Chr 8, 66523076 |
| Off-target 2 | GTCATGTGACAGATGGGGCAGGGG, Chr 5, 149800667 |
| Off-target 3 | GTCTGAAGACAGATGGGGCCCAGG, Chr 5, 6336730 |
| Off-target 4 | GTCTGTTGACAGTTGGGGCAGTGG, Chr 5, 175712652 |
| Off-target 5 | GTCTGTGCACACATCCGCCCGCGG, Chr 5, 181182354 |
| **gRNA2 Off-targets** | |
| Off-target 1 | CGGCGCTCGGGCTCAGCTGTCCGG, Chr 8, 12951655 |
| Off-target 2 | CTGCATCAGGCTTAACATGACTGG, Chr 20, 46237257 |
| Off-target 3 | CTACTTAGGGCTGCAACTGACTGG, Chr 1, 59950252 |
| Off-target 4 | CTGTGTCAGGGCTGAACTGAGGGG, Chr 22, 49642616 |
| Off-target 5 | GTGCCTCGGGCTCAGACTGACCGG, Chr 7, 159033388 |
| **AAVS1 gRNA Off-targets** | |
| Off-target 1 | GGAGACATTAGGGACAGGATAAG, Chr 10, 119439168 |
| Off-target 2 | GGGACCATCAGGGACAGGATGGG, Chr 6, 36797686 |
| Off-target 3 | GAGGGCAGCAGGGACAGGAGGG, Chr 12, 131827311 |
| Off-target 4 | GAGGACAGTAGGGACAGGTTAAG, Chr 18, 8749293 |
| Off-target 5 | GGGTGCAGTGGGGACAGGATGGG, Chr 20, 57806974 |
